# Cooling fast and slow: Characterising the effects of vitrification in cryo-EM and the subsequent recovery of equilibrium populations

**DOI:** 10.64898/2026.04.21.720011

**Authors:** Robert Clark, Louis G. Smith, Matthew P. Leighton, Ryan Szukalo, Syma Khalid, Pablo G. Debenedetti, Pilar Cossio, Miro A. Astore, Sonya M. Hanson

**Affiliations:** Department of Biochemistry, University of Oxford, Oxford, UK; University of Pennsylvania Department of Biochemistry and Biophysics, Philadelphia, PA, USA; Department of Chemistry, Princeton University, Princeton, NJ, USA; Department of Physics and Quantitative Biology Institute, Yale University, New Haven, CT, USA; Center for Computational Biology, Flatiron Institute, New York, NY, USA; Center for Computational Mathematics, Flatiron Institute, New York, NY, USA; Department of Chemical and Biological Engineering, Princeton University, Princeton, NJ, USA

**Author notes:** These authors contributed equally to this work (Robert Clark, Louis G. Smith).

## Abstract

Single-particle cryogenic electron microscopy (cryo-EM) has enabled near-atomic resolution structure determination of diverse biomolecules. Because the high vacuum required for electron microscopy prevents the imaging of liquid-phase samples, cryo-EM samples are prepared by plunging the sample into a cryogen, rapidly cooling the sample and suspending the ensemble of biomolecules in a matrix of water glass. However, the effects of this vitrification on the biomolecular ensemble are unknown, complicating efforts to use cryo-EM to derive conformational ensembles of biomolecules. To study these effects, we carried out extensive molecular dynamics simulations (over 50 milliseconds) of the Trp-cage miniprotein at equilibrium and undergoing rapid cooling. We simulated seven cooling rates spanning three orders of magnitude, with the slowest coolings matching experimental rates. By inspecting molecular mobility and density-temperature equations of state for water with and without protein, we found that water vitrification is unaltered by the protein. To track protein conformation changes, and to relate them to conformational kinetics, we made a Markov State Model (MSM) of Trp-cage from 5.4 milliseconds of equilibrium sampling at 277 K. We observed that MSM states with a characteristic time longer than the duration of the non-equilibrium cooling, tend to be more robust to artefacts induced by such cooling. Critically, although we observe that some states vanish in the equilibrium ensemble at 230 K, none do in our nonequilibrium cooled ensembles. However, to account for perturbations induced by nonequilibrium cooling for more labile states, we developed a thermodynamic inference framework for recovering equilibrium populations from the measured vitrified ensembles. These results indicate that cryo-EM has the capacity to be a reliable and accurate biophysical technique for the study of biomolecular ensembles.

**Significance:** Cryogenic electron microscopy images biomolecules trapped in vitreous ice. To vitrify the sample, it must be cooled over the course of 22 microseconds. However, the degree to which this cooling causes the ensemble of the molecules to be perturbed from equilibrium is unknown. Here we present extensive molecular dynamics simulations to quantify the equilibrium dynamics of the Trp-cage miniprotein and the effects of cooling on its conformational ensemble. By simulating cooling at seven different rates, including the slowest experimental rates that still result in vitrification, we connect the kinetic properties of a protein’s conformational state to the change in state population from cooling. We show that cooling-induced population shifts are small but observable. We further introduce a thermodynamic-inference method to recover equilibrium populations from the cooled ensembles.

## Introduction

Elucidating the conformational changes biomolecules undergo is fundamental to understanding their function. Proteins, nucleic acids, and their complexes exist as dynamic ensembles, sampling multiple conformations that enable binding, catalysis, and regulation [1]. Cryogenic electron microscopy (cryo-EM) is a transformative tool for structural biology, enabling near-atomic resolution structure determination of biomolecules without the need for crystallization [2]. Single-particle analysis routinely resolves multiple distinct conformational states from the same sample [3], and emerging computational methods now aim to characterise continuous conformational landscapes and state populations [4, 5]. It is often assumed that the ensemble imaged in cryo-EM resembles the equilibrium ensemble of the sample right before vitrification, however the veracity of this assumption remains an open question [6, 7], limiting the insights one can make from biomolecular conformations observed in cryo-EM data.

The high vacuum required for electron microscopy prevents the use of liquid-phase samples, and so cryo-EM relies on plunge-vitrification into a cryogen, usually liquid ethane, for sample preparation. This technique cools the sample to below 90 K within microseconds [8, 9] (Fig. 1A). The rapid cooling prevents the formation of hexagonal ice, which strongly scatters electrons, disrupting imaging, and structurally damages the biomolecules [2]. Instead water molecules arrest into low-density amorphous ice (LDA) [10, 11], a metastable glass that suspends macromolecules in a rigid, electron-transparent matrix [12]. In 1988 Jaques Dubochet, one of the inventors of the vitrification process, published an estimate for the minimum cooling rate required to successfully vitrify water. By extrapolating from measurements of the fastest achievable cooling rate for thin specimens of tissue, Dubochet estimated that a cooling rate of 10^6^ K/s could be achieved for a 1 µm thick film of water [2]. This set the order of magnitude estimate for the critical cooling rate of water until the invention of an apparatus that allowed careful control of the vitrification process via laser mediated heating. With this more advanced instrumentation, the minimum cooling rate for vitrification has now been precisely measured at 6.4 × 10^6^ K/s, a number in agreement with Dubochet’s estimate nearly 40 years earlier [9]. At this cooling rate, cooling from 4^°^C (∼ 277 K) to past the glass transition (*T*_*g*_) point at ∼136 K occurs over approximately 22 µs, a time scale that overlaps with many biomolecular conformational transitions and binding events [13].

**Fig. 1.**
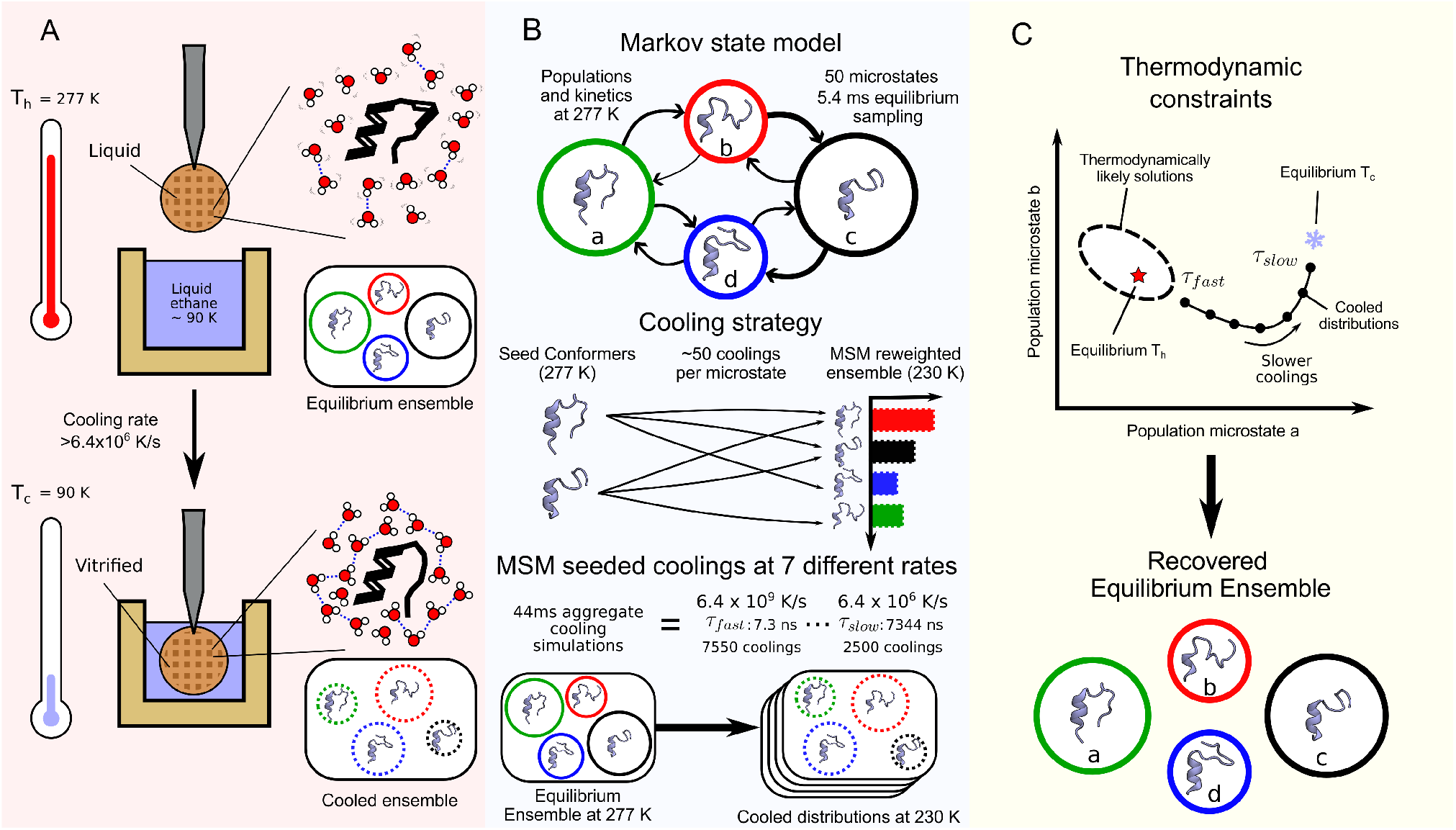
Investigating and recovering from the effects of rapid vitrification on biomolecular conformations. **A.** Cryo-EM relies on vitrification of solvated biomolecules to withstand high vacuum, immobilise, and preserve them. Samples are blotted onto a grid and pluged into liquid ethane, causing the solvent to transition into low density amorphous water (LDA). **B**. The equilibrium ensemble at several temperatures was sampled through temperature replica exchange simulations, and the kinetics of the system at 277 K were estimated using a Markov State Model (MSM) built from equilibrium MD trajectories. To obtain vitrified ensembles for different cooling rates, we cooled conformers from each MSM state roughly 50 times at seven different cooling rates, resulting in tens of thousands of cooling trajectories from 277 K to 230 K. By recording the MSM state identity of conformations before and after cooling. **C**. Thermodynamic constraints provide bounds on the maximum perturbations that nonequilibrium cooling can have on the equilibrium ensemble. By combining a series of these cooled ensembles, we found that these constraints are sufficiently narrow to enable accurate recovery of an equilibrium ensemble from the nonequilibrium cooled ensembles, in a process called thermodynamic inference.

Conformational transitions with time scales comparable to the cooling duration could occur during vitrification, allowing populations to evolve away from the initial 277 K ensemble. Indeed, given sufficiently slow cooling, we would expect the ensemble to relax to the low temperature ensemble. Equilibrium conformational populations are known to shift at low temperature, with some proteins undergoing cold denaturation [14]. At the other extreme, kinetic trapping could occur, where thermal fluctuations are not sufficient to permit equilibration of the system at all, which would retain the original equilibrium present prior to cooling. The extent of thermal relaxation possible during vitrification ought to determine the degree of resemblance between vitrified ensembles and those at 277 K. One can illustrate the complicated balance between kinetic trapping and thermal relaxation even using a simple 1D toy model (Fig. S1). Simulating Langevin dynamics on a simple four well potential undergoing rapid cooling shows how different barrier heights are overcome at different cooling rates. At fast rates, cooled ensembles still resemble the hot equilibrium, and are thus fully kinetically trapped, whereas for slow cooling the ensemble matches the cold equilibrium, since it has time to relax. We will refer to such relaxation as annealing in what follows.

At intermediate cooling rates, the outcome is not a simple interpolation between these limits. Instead, the competition between kinetic trapping and annealing results in states that are enriched at one rate and depleted at another, while more stable states remain unchanged (Fig. S1B). The complexity of outcomes seen even in this simple model motivates a full investigation of how a protein ensemble evolves under cooling at realistic cryo-EM cooling rates, and which properties make particular states more susceptible to change during vitrification. This is especially important given experimental evidence that the temperature prior to cooling influences the conformational distributions observed in cryo-EM [15, 16], consistent with both kinetic trapping or partial thermal relaxation during cooling, and in light of emerging time-resolved cryo-EM experiment where the achievable temporal resolution is approaching the vitrification time scale [17].

Computational methods are well-positioned to address how the balance of kinetic trapping and thermal relaxation changes the conformational distribution of a biomolecule during vitrification. For example, Bock and Grubmüller [18] performed molecular dynamics (MD) simulations of ribosome vitrification at cooling rates of 10^9^ to 10^11^ K/s, demonstrating that energy barriers below approximately 10 kJ/mol can be traversed during cooling. However, these simulations used cooling rates 1000-fold faster than the fastest experimental rate, and analysis was based on a two-state framing. Given our more detailed experimental understanding of the vitrification process and recent results in stochastic thermodynamics that use constraints on entropy production in nonequilibrium processes for inference [19, 20], we can also now apply these approaches to the nonequilibrium cooling process that occurs in cryo-EM.

We found the need to move away from a characterisation of the kinetics as emerging from a lone relevant barrier. Instead, to score the connectedness of a state in the kinetic network, we borrowed from existing stochastic thermodynamics literature on the dynamical activity in discrete-state Markov models [21] and define a characteristic time over which a given state exchanges probability with the rest of the network. We name it the *event time* of a MSM state *k*, and express it as *v*_*k*_ (τ) ≡ 0.05 × 2τ*/a*_*k*_ (τ) (for more discussion see Eq. (1) and also the Methods section ‘Categorizing change under cooling with hot equilibrium kinetics’). Here, τ is the lagtime, and *a*_*k*_ (τ) is the part of the dynamical activity contributed by state *k* at that lagtime. The factor of 0.05 scales the quantity to be the amount of time needed to flow 5% of the system’s total population. The event time therefore gives the mean time required for 5% of the system’s population to be exchanged to other states. This is a measure of gross probability traffic associated to a particular state.

Here we combined extensive molecular simulation and approaches in stochastic thermodynamics to investigate perturbations to conformational ensembles introduced by vitrifaction and then recover the equilibrium distribution from the the vitrified ensembles. Our system of choice is the Trp-cage miniprotein. Its small size (20 residues) permits exhaustive conformational sampling while exhibiting rich dynamics including microsecond time scale folding [22, 23] and cold denaturation [14, 24]. We used the TIP4P/ICE water model, due to its accuracy near the freezing point [25, 26]. We characterised the equilibrium landscape of Trp-cage at multiple temperatures and built a Markov state model (MSM) to describe the kinetics at 277 K. We ran tens of thousands of nonequilibrium cooling trajectories with conformations drawn from each microstate at experimentally realistic cooling rates (Fig. 1B) [9]. By tracking the occupancy of MSM states before and after cooling, we analyse how a state’s equilibrium kinetics influence its population changes under nonequilibrium cooling, and infer the general features of protein states beyond Trp-cage that influence behaviour under cooling. We find that the population of a state may drift during cooling when the cooling time exceeds the event time, defined above. Finally, we developed a framework for thermodynamic inference using constraints on entropy production during cooling to recover the original 277 K equilibrium distribution from the cooled distributions (Fig. 1C). Overall, these findings provide a robust framework to understand when states are disrupted by vitrification and how to correct for these cooling artifacts, both critical insights to the accurate interpretation of biomolecular ensembles recovered from cryo-EM samples.

## Results

### Hot and cold equilibrium properties of the Trp-cage system

To understand how rapid cooling affects biomolecular ensembles, we first characterised Trp-cage’s properties at equilibrium. We performed temperature replica exchange (TREX) MD simulations spanning 230 K to 520 K using 90 replicas with 12 µs of sampling per replica for a total of 1.08 ms. We chose 230 K as our lowest temperature because in pilot nonequilibrium cooling simulations this temperature marked the onset of vitrification (Fig. S2), where both conformational changes and water self-diffusion slow dramatically (Fig. S3). The TREX revealed markedly different ensembles at 277 K versus 230 K (Fig. 2A, Fig. S4). In what follows, we use ‘microstate’ and ‘state’ interchangeably to refer to the states of our Markov model.

**Fig. 2.**
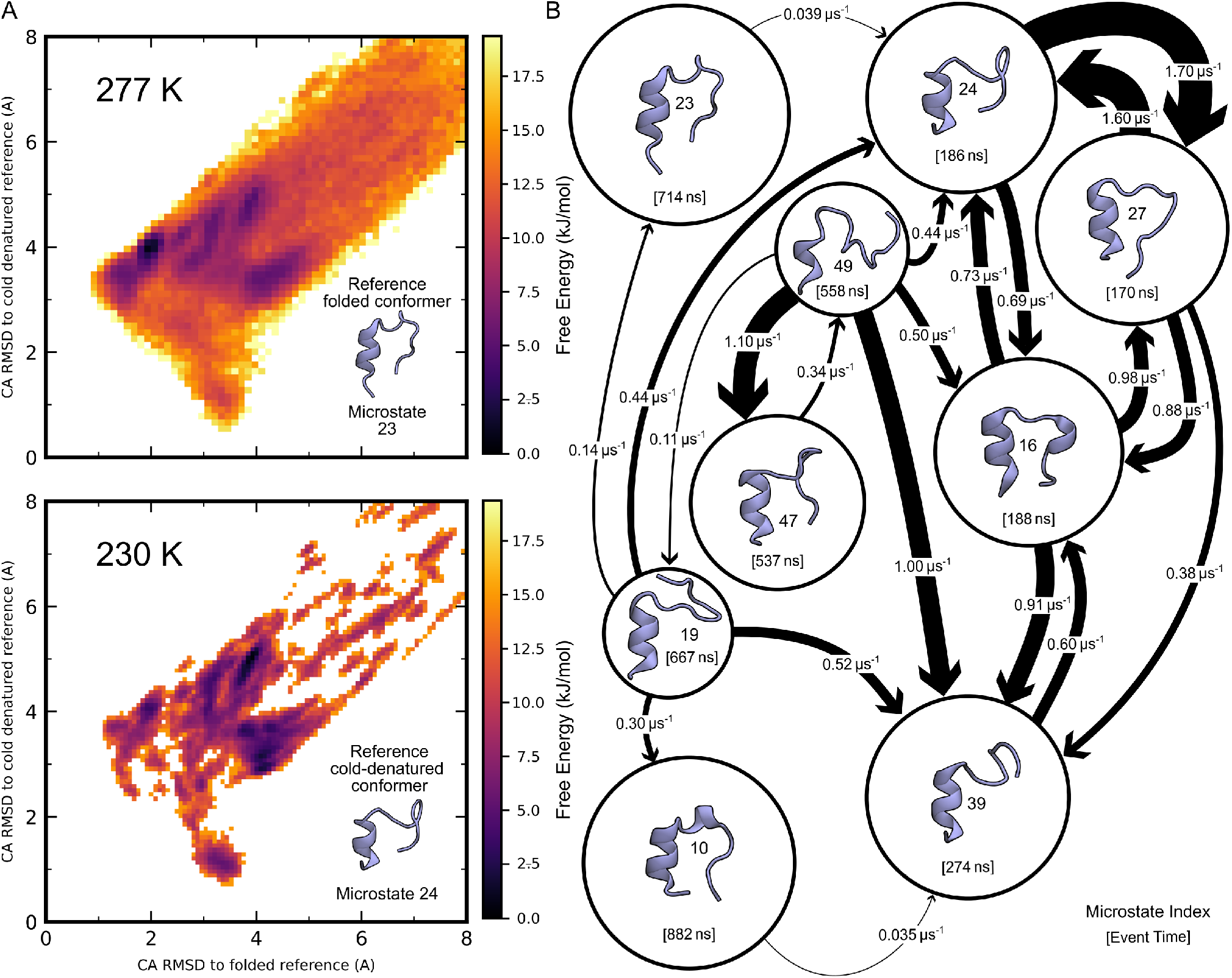
Markov state model with 50 microstates describing the equilibrium conformational landscape of Trp-cage at 277 K and 230 K. **A.** Free energy surfaces of Trp-cage at 277 K (top) and 230 K (bottom) from temperature replica exchange MD simulations. The coordinate axes show C_α_ RMSD to cold-denatured reference (y-axis) and C_α_ RMSD to folded reference (x-axis). Reference structures are shown inset; see Fig. S6 for cluster centre location on landscape. **B**. Markov state model connectivity graph showing transition rates between the nine states with probabilities greater than 1% at 277 K. Circle diameter is proportional to the log state population and the event time associated with each microstate is in square brackets. Arrow thickness is linearly proportional to the transition rate between states, with values labelled in µs^−1^. Only transitions with rates *>* 0.3 µs^−1^ are shown for visual clarity, except in cases where a rate below the cutoff was included to demonstrate that every state has in and out transitions.

At 277 K, Trp-cage predominantly populates the native folded conformations (states 10 and 23), consistent with experimental characterisation [23]. At 230 K, Trp-cage favours multiple cold-denatured conformations where the *α*-helix or the 3_10_-helix begin to lose their structure (states 27, 39, 47 and 49). Additionally, there are ten microstates in the 277 K ensemble which we do not sample in the 230 K ensemble (Fig. S4). These temperature-dependent equilibrium shifts serve as fiducials for evaluating vitrification effects. That is, if the cooling were slow enough to allow full thermal relaxation, we would find that the populations after cooling match those from the equilibrium distribution at 230 K. Therefore, deviations from this distribution reflect the particulars of nonequilibrium cooling.

To quantify kinetics at 277 K and track conformational changes during cooling, we performed 5.4 ms of unbiased Trp-cage simulations with conformations seeded from across the 277 K TREX ensemble. We clustered and featurized trajectories using all rotatable dihedral angles reduced with time-lagged Independent Component Analysis (TICA) [27, 28]. From these we built a Bayesian MSM with 50 microstates using a 40 ns lag time, and constrained the stationary distribution to the 277 K populations from TREX (see Methods) [29]. As the simulations were performed in explicit solvent, the model implicitly reflects solvated conformational preferences. The resulting kinetic network reveals the connectivity and transition rates between states (Fig. 2B, Fig. S5). The MSM showed clear kinetic separation between different cold-denatured conformations and multiple similar but distinct folded states (Fig. S6). The range of timescales provided a suitable range to relate the rate of cooling up to and including experimentally relevant states. To compare with the non-equilibrium cooling dynamics, we begin by asking whether water’s behaviour is perturbed in the protein and water system cooled using our sampling approach.

### Bulk solvent vitrification is unperturbed by protein presence

To study the bulk solvent under cooling, we characterised water vitrification through molecular mobility and density-temperature equations of state. We found that neither the presence of the protein nor the cooling rate had a significant effect on the thermodynamic pathway or dynamics of vitrification, which proceed as in pure bulk water. Simulation snapshots where water molecules are colour-coded by their displacement over 0.5 ps illustrate the progressive slowing of water dynamics upon cooling (Fig. 3A). At 277 K, water molecules exhibit large, thermally-driven displacements typical of a normal liquid. By 230 K, mobility has decreased substantially and dynamic heterogeneity emerges, with pockets of mobile water molecules coexisting alongside near-arrested ones, behaviour characteristic of deeply supercooled liquids approaching vitrification. At 100 K, nearly all water molecules are immobile, with displacements reduced to small-amplitude vibrations characteristic of glassy dynamics. Velocity autocorrelation functions further characterize this dynamical transition, showing increasingly oscillatory molecular motion upon cooling, and reveal that only the first solvation shell exhibits dynamics distinct from bulk water, with the degree of this perturbation remaining largely independent of temperature (analysis not shown).

**Fig. 3.**
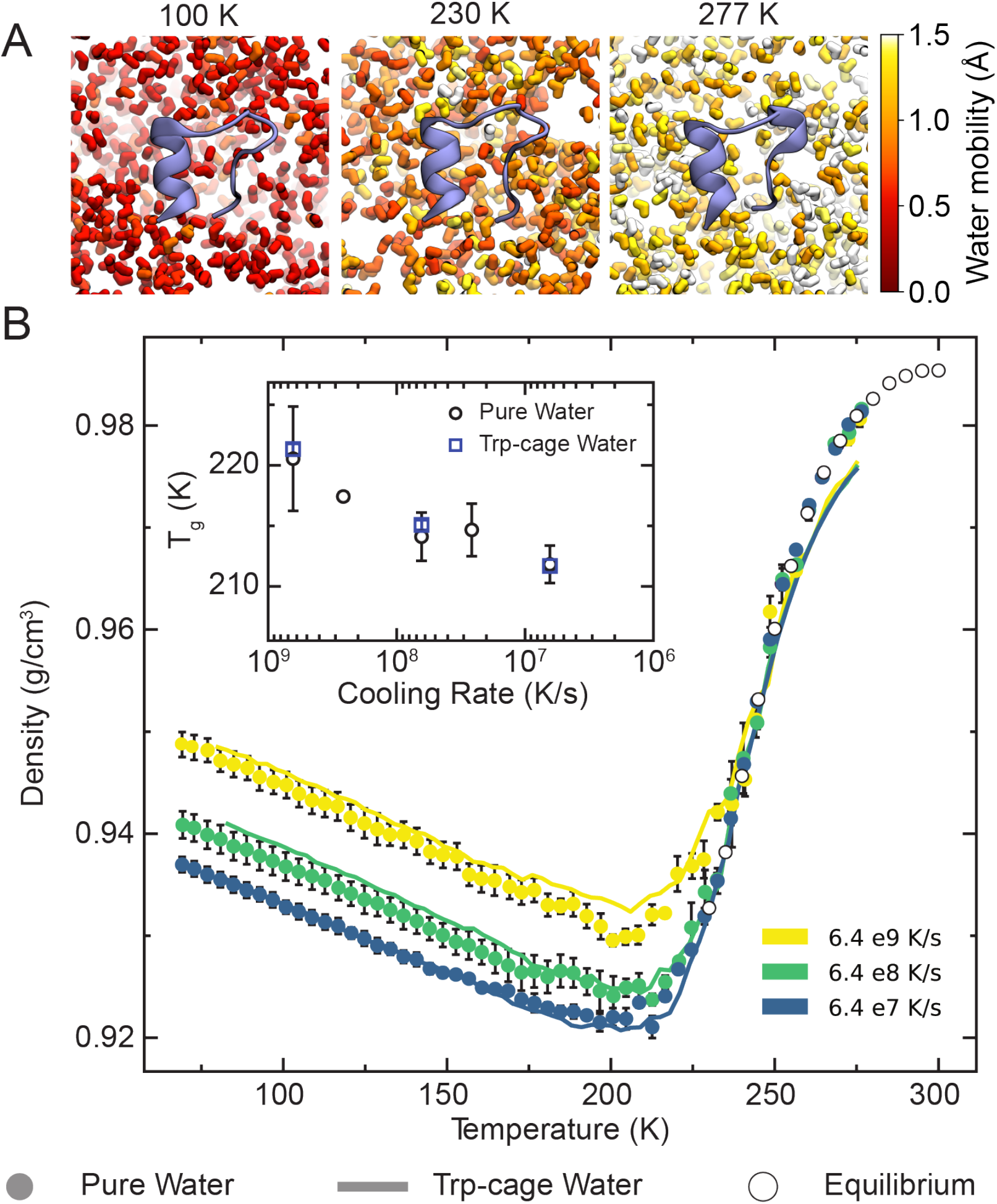
Water vitrification dynamics and thermodynamics with and without protein. **A.** Snapshots of water molecules coloured by displacement over 0.5 ps at three temperatures (100 K, 230 K, 277 K), showing progressive reduction in water mobility during cooling. Red indicates low mobility, white indicates high mobility. **C**. Water density as a function of temperature during isobaric cooling at three different cooling rates. Filled circles represent pure water systems, while solid lines show water density in Trp-cage-containing systems (excluding protein volume). Open circles indicate equilibrium density values from NPT simulations. Colours distinguish cooling rates: 6.4 *×* 10^9^ K/s (yellow), 6.4 *×* 10^8^ K/s (green), and 6.4 *×* 10^7^ K/s (blue). **Inset:** Glass transition temperature, *T*_g_, as a function of cooling rate for pure water (circles) and Trp-cage systems (squares). Error bars represent standard error of the mean across independent simulations.

To assess the thermodynamic pathway during cooling, we compared rate-dependent density-temperature equations of state for pure water and water in the presence of Trp-cage (Fig. 3B), extending cooling simulations to 80 K to fully characterize the glassy regime. For the Trp-cage simulations, the effective water density was computed by excluding the protein volume, taken as the volume occupied by spheres defined by the atomic van der Waals radii, from the simulation cell volume (Fig. S7). At high temperatures, all cooling rates reproduce the equilibrium NPT density (open circles), confirming that the simulations are well-equilibrated at the onset of cooling. As temperature decreases, slower cooling rates track the equilibrium curve to lower temperatures before diverging, indicating that the systems remains in equilibrium longer at slower rates. In all cases, the density reaches a minimum near 215 K before increasing as the system enters a glassy state, hallmarks of water’s anomalous behaviour and the formation of LDA [11]. Critically, the density-temperature curves for Trp-cage-containing water are indistinguishable from pure water at all three cooling rates, with neither the density minimum nor the final glassy-state density affected by the miniprotein’s presence.

Further, we calculated *T*_*g*_ for both pure water and the Trp-cage system at each cooling rate using a global hyperbolic fitting procedure (Fig. 3, inset) [30, 31]. As expected, *T*_*g*_ decreased monotonically with decreasing cooling rate. Importantly, the *T*_*g*_ values for Trp-cage simulations matched pure water at each cooling rate, showing that a single Trp-cage molecule does not measurably alter water vitrification.

These results establish that the presence of a small, compact biomolecule does not perturb either the thermodynamic pathway or the glassy dynamics of supercooled water during rapid cooling, and validate our choice of 230 K as the cooling endpoint for protein conformational analysis. By this temperature, water mobility has decreased dramatically and the system approaches the glassy regime where conformational changes effectively cease. Having established that water vitrification proceeds normally in the presence of protein, we can proceed with investigations of the protein conformations as a function of cooling.

### The influence of cooling on the conformational ensemble of a macromolecule

Having characterised the equilibrium landscape and kinetics of the Trp-cage system, we employed a shooting strategy to sample the effect of cooling on the conformational ensemble at seven cooling rates spanning three orders of magnitude, from 6.4 × 10^9^ K/s (duration of 7.3 ns) up to the critical cooling rate of 6.4 × 10^6^ K/s (duration of 7.3 µs) [9], accumulating 44 ms of sampling (Supplementary Table 1). For each cooling rate, we started collections of simulations from conformers obtained by geometric clustering of the TREX samples (see Methods).

One might suppose that perturbations in protein state populations induced by cooling could yield an ensemble similar to an equilibrium ensemble at some temperature between the starting hot temperature and finishing cold temperature. To test this, we computed the Kullback-Leibler (KL) divergence between equilibrium populations at each temperature from the TREX simulations and the distributions resulting from cooling (cooled distributions) (Fig. 4A). For all cooling rates, the equilibrium distribution at 277 K remains the closest match to the cooled ensemble, with KL divergence increasing monotonically for lower temperatures.

**Fig. 4.**
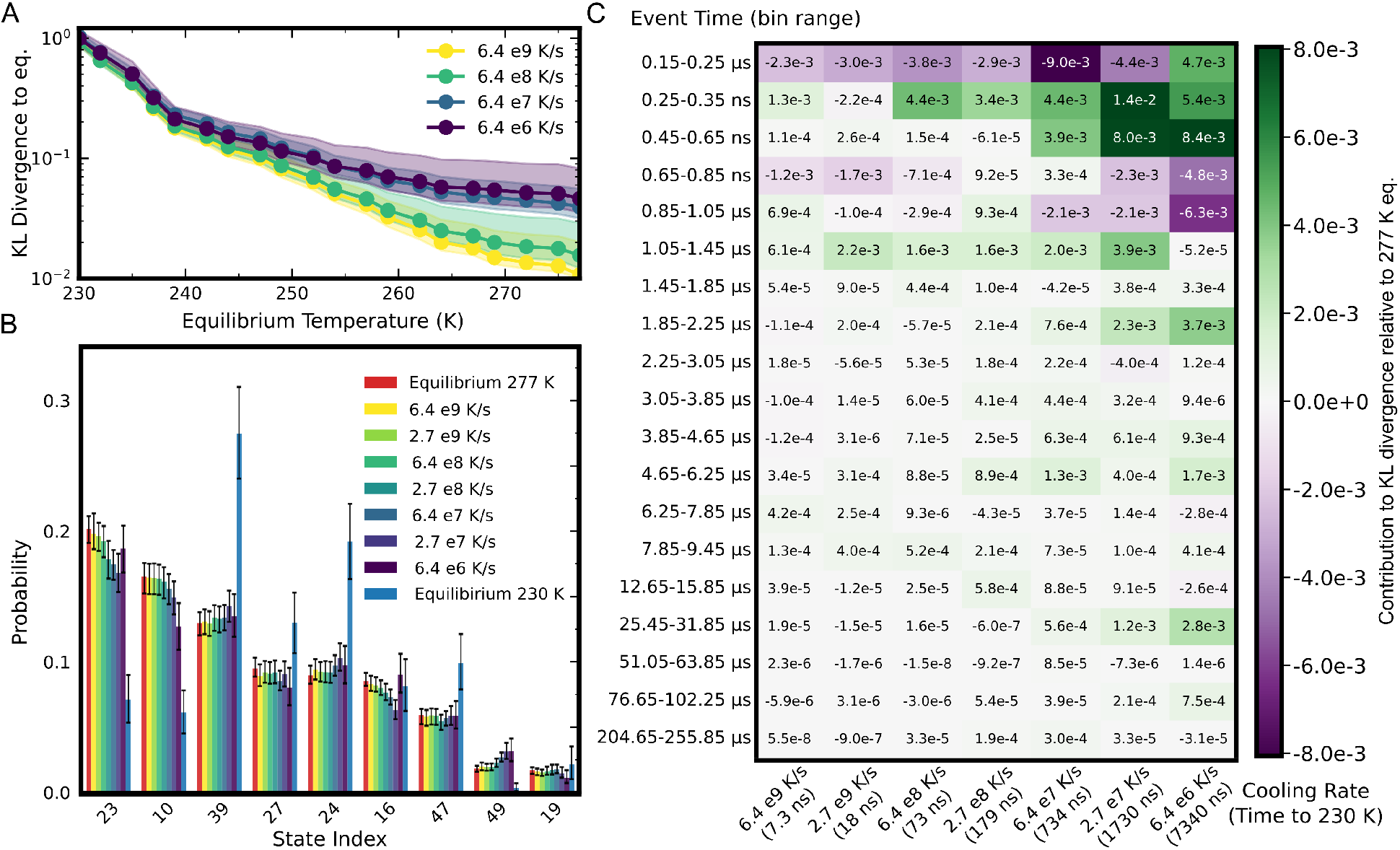
Cooling rate-dependent changes to the conformational distribution of Trp-cage. **A.** Kullback-Liebler (KL) divergence between cooled population distributions and equilibrium population distributions at each temperature from 277 K to 230 K, using the equilibrium distribution at each temperature as a reference. Each line represents a different cooling rate and shaded area represents 95% credible intervals. **B**. State populations following vitrification at multiple cooling rates (colored bars) compared to equilibrium state populations at 277 K (red bars) and 230 K (blue bars). States with population *>*1% at 277 K are shown. All 50 states can be found in Fig. S8A. Error bars represent 95% credible intervals from Bayesian bootstrapping. **C**. Heat map showing the relationship between states’ event times, the cooling rate, and median change in that bin’s contribution to the KL divergence between the equilibrium 277 K and cooled distributions, with the equilibrium as the reference. Numbers in the heat-map also report this value. States are binned together by their event times. See Fig. S8B for a per-state heat map.

Cooling induces modest but measurable population shifts in the most populous states (Fig. 4B). At the fastest cooling rate (6.4×10^9^ K/s), populations remain indistinguishable from their 277 K equilibrium values for all nine states with abundance greater than 1%. As cooling slows, certain states exhibit systematic deviations. For instance, state 10, a folded state comprising 16.5% of the 277 K equilibrium, decreases to 12.7% at the critical cooling rate. However, in some cases states do not move toward their equilibrium population at 230 K (e.g state 49). We also observed non-monotonic population change (for example, Fig. S8A states 8, 38, 46). This is consistent with the behaviours of our toy model (Fig. S1). Despite these changes, all states remain populated even at the slowest cooling rate. Notably, within our state-space, there are ten zero population states at equilibrium at 230 K (Fig. S4). This result is critical for efforts to reweight populations obtained from cryo-EM data, as absent conformations become unknowns.

We hypothesized that kinetic network surrounding a state may categorize the extent of perturbation. To quantify this kinetic network, we defined an “event time” for each microstate (See Methods section ‘Categorizing changes under cooling with hot equilibrium kinetics’),

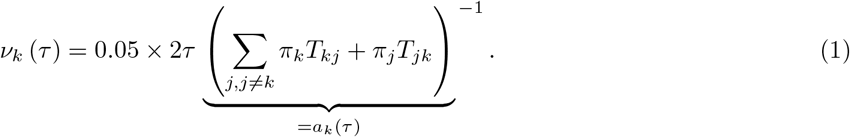

Here, *v*_*k*_ is the event time for state *k*, τ is the lagtime, *a*_*k*_ (τ) is the part of the dynamical activity contributed by state *k* at that lagtime, *π*_*j*_ is the population of state *j*, and *T*_*jk*_ is the transition probability for going from *j* to *k* with lagtime τ, and the factor of 0.05 scales the quantity to be the amount of time needed to flow 5% of the system’s total population. Broadly, we expect that states with slowest event times will be resistant to annealing. Note that almost all states with long event times are so kinetically isolated that they are estimated to be rare at equilibrium (Fig. S8A). In our system, the states with the fastest event times are 24 and 27 (186 and 170 ns respectively), while the slowest event time (205 µs) is exhibited by state 31 (Fig. S8B). We binned states according to their event time (Eq. 1) and calculated the per state contribution to the KL divergence (See Methods) from the 277 K equilibrium distribution as a function of the cooling rate. At the fastest cooling rate, states exhibited almost no changes in population. As cooling slows, states with event times comparable to or shorter than the cooling duration show population changes, whereas states with event times exceeding the cooling time do not.

### Changes to protein observables from vitrification

Having characterised ensemble-level perturbations, we investigated how cooling affects protein structure. To probe structural changes beyond the resolution of our MSM (limited by its 40 ns lag time), we examined sidechain dihedral angles, which can undergo rotameric transitions on faster time scales. We focused on two representative residues: Asn1, a solvent-exposed asparagine at the N-terminus and Trp6, the tryptophan that forms the hydrophobic core. The *χ*_1_ distribution of Asn1 shows clear cooling-rate dependence (Fig. 5A). At 277 K equilibrium, this sidechain primarily samples three rotameric states. After cooling, the distribution shifts toward gauche (+) with slower cooling rates, corresponding to the rotamer most favoured at lower temperatures. In contrast, the χ_1_ distribution of Trp6 remains essentially unchanged between hot equilibrium and all cooled ensembles (Fig. 5B). More generally, solvent-exposed residues tend to show cooling-rate-dependent sidechain reorientations, while buried residues in the hydrophobic core remain largely unaffected (Figs. S9 and S10). Consistent with this, the ensembles we sample tend to be more compact, in terms of their *R*_*g*_, at the slowest cooling rate we consider, but not as compact as the equilibrium 230 K limit (Fig. 5C).

**Fig. 5.**
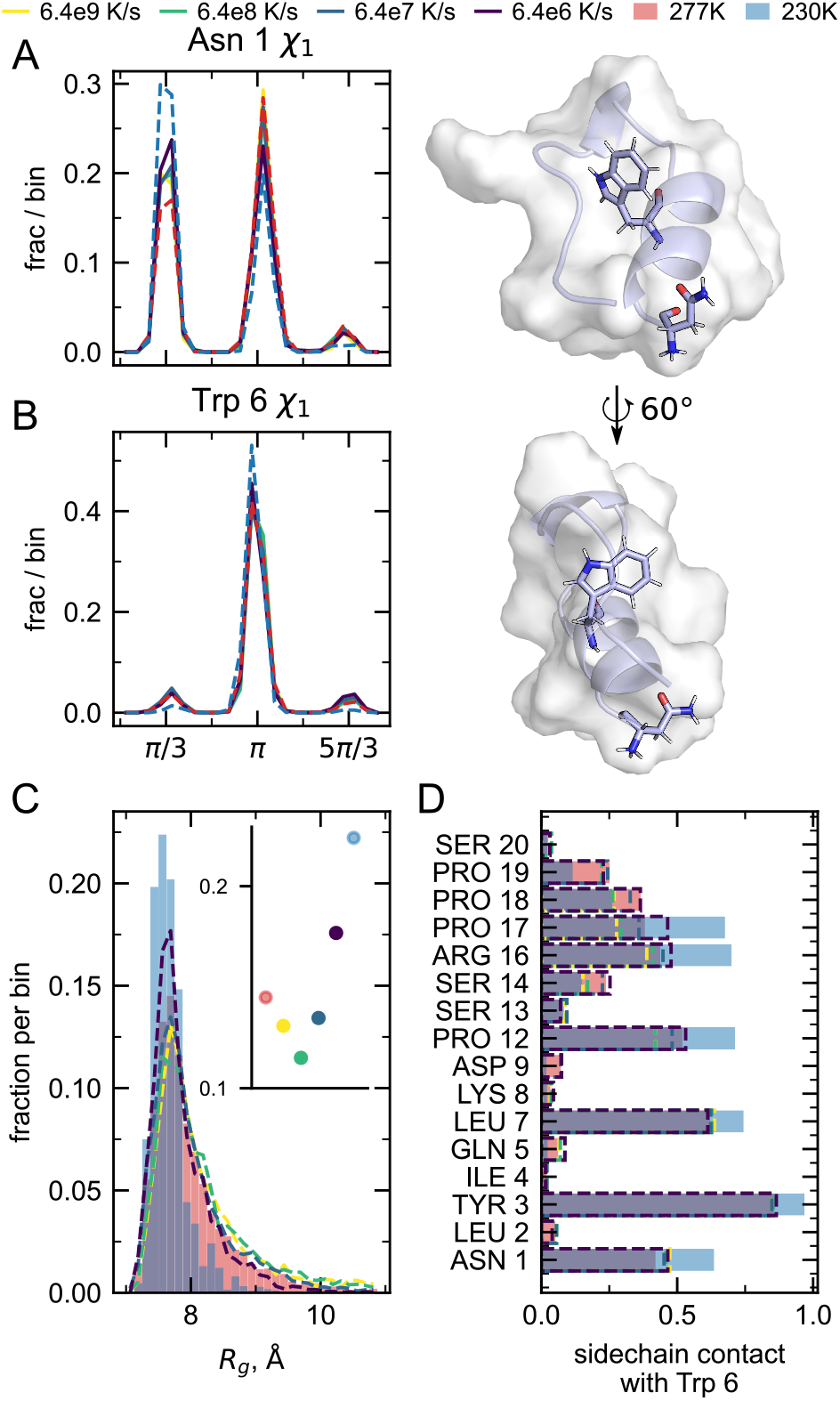
Structural changes during vitrification localize to solvent-exposed regions while preserving the hydrophobic core. **A, B.** Sidechain *v*_1_ dihedral angle distributions at equilibrium (277 K, red, and 230 K, blue) and after cooling at multiple rates (dashed lines) for (A) Asparagine 1, a solvent-exposed residue, and (B) Tryptophan 6, a buried core residue (both are shown in renders to the right). Angles are shifted by τ for easier visualization. **C**. Radius of gyration (*R*_*g*_) distributions. Filled red bars show 277 K equilibrium; filled blue bars show 230 K equilibrium; coloured lines show cooled distributions. The inset shows the maximum height of each histogram, on the Y axis. **D**. Fraction of contact between each other sidechain in the protein and Trp6’s sidechain, where contact is defined as any heavy atom within 4.5 Å. Colours in all panels match the legend.

Despite these global changes in compaction, the protein’s hydrophobic packing remains largely intact. We quantified core structure by measuring sidechain-sidechain contacts between Trp6 and all other residues (Fig. 5E). The pattern of contacts in cooled ensembles closely matches the 277 K equilibrium across all cooling rates. Residues that make frequent contacts with Trp6 at 277 K, such as Pro12, Tyr3, and Pro18, maintain similar contact frequencies after cooling, while residues that rarely contact Trp6 at equilibrium remain unassociated in cooled ensembles.

Taken together, these structural analyses reveal that vitrification-induced changes concentrate in the flexible, solvent-exposed periphery of the protein. Slower cooling allows partial compaction and sidechain rearrangements in exposed regions as the system begins to relax toward more compact configurations accessible at reduced temperatures. However, these changes do not generally correspond to adoption of a different overall protein architecture, with the fold and hydrophobic core remaining broadly preserved, even while state populations are being affected by vitrification.

### Thermodynamic constraints enable the recovery of equilibrium populations from cooled ensembles

Given we observed perturbed state populations resulting from vitrification, this motivated the development of a method to recover the equilibrium ensemble from a collection of cooled ensembles. Here, we developed an algorithm for thermodynamic inference based on bounds for entropy production derived from the principles of stochastic thermodynamics [19, 20]. These thermodynamic constraints allowed us to determine the accessible equilibrium ensembles from the distributions at multiple cooling rates.

We studied the thermodynamics of nonequilibrium cooling by considering two distinct processes: the initial cooling, from time *t* = 0 at temperature *T*_*h*_ = 277 K to time *t* = τ at temperature *T* = 230 K, and an auxiliary relaxation at temperature *T*_*c*_ = 230 K from *t* = τ onward. At time *t* = 0 the ensemble is the equilibrium Boltzmann distribution *π*(*T*_*h*_) at temperature *T*_*h*_, while in the long time limit as *t* → ∞ the ensemble will asymptotically approach the equilibrium Boltzmann distribution *π*(*T*_*c*_).

We modelled the cooled protein using master equation dynamics on a set of discrete states corresponding to the microstate MSM, each assumed to be characterised by state-dependent energy and entropy *e*_*i*_ and *s*_*i*_. The state entropies account for the fact that the MSM microstates are not true microstates from a statistical mechanics standpoint [32]. We note that our having these entropies implies that the MSM microstates occupy a range of volumes in phase space. We do not explicitly account for interactions between the protein and the solvent, however the state-dependent energies and entropies can implicitly account for such interactions.

We built on known results from stochastic thermodynamics [33] to compute the entropy production for the two processes under consideration. For each cooling time τ, the entropy productions Σ_cool_ and Σ_aux_ of the initial cooling and auxiliary relaxation processes, respectively, are functions of the state-dependent energies and entropies (both unknown), and the state probabilities at the end of the cooling process *p*(τ) (measured from experiments). We then applied the second law, requiring that entropy production of both the initial cooling process, as well as the auxiliary relaxation process, must be non-negative for all cooling times τ :

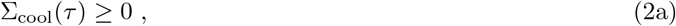

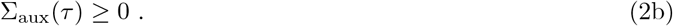

We additionally took advantage of the data from multiple distinct cooling times by considering the relationships between entropy productions of cooling and relaxation for adjacent cooling times. We conjectured that the entropy production of the cooling process increases monotonically with cooling time, since for slower cooling rates the protein ensemble has more time to change and thus dissipate heat to its surroundings. Conversely, we conjectured that the entropy production of the auxiliary relaxation process decreases monotonically with cooling time, as the slower the cooling process the closer the ensemble should be to reaching thermodynamic equilibrium at *T* = 230 K. By requiring monotonicity of entropy production, we obtained two additional inequality constraints for each pair of adjacent cooling times, thus taking advantage of the multiple cooling rates considered in this study. Specifically, for a set of *n* cooling times τ_1_ *<* τ_2_ *<* … *<* τ_*n*_, we enforced

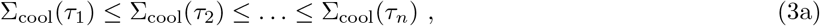

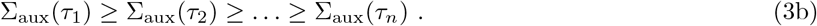

Details for calculating entropy production are in Methods section ‘Thermodynamic constraints on cooling’.

We then used the thermodynamic constraints to recover the initial equilibrium distributions, with an algorithm implemented as follows: first, we sampled points in the space of state entropies and valid probability distributions (satisfying normalization), then iterated a differential evolution algorithm until all thermodynamic inequalities were satisfied. Repeating this process for many randomly-drawn initial points allowed us to sample the space of thermodynamically-allowed distributions. This ensemble of allowed distributions is displayed through white shading in Fig. 6A, which shows a projection of the state distribution onto two particular states.

**Fig. 6.**
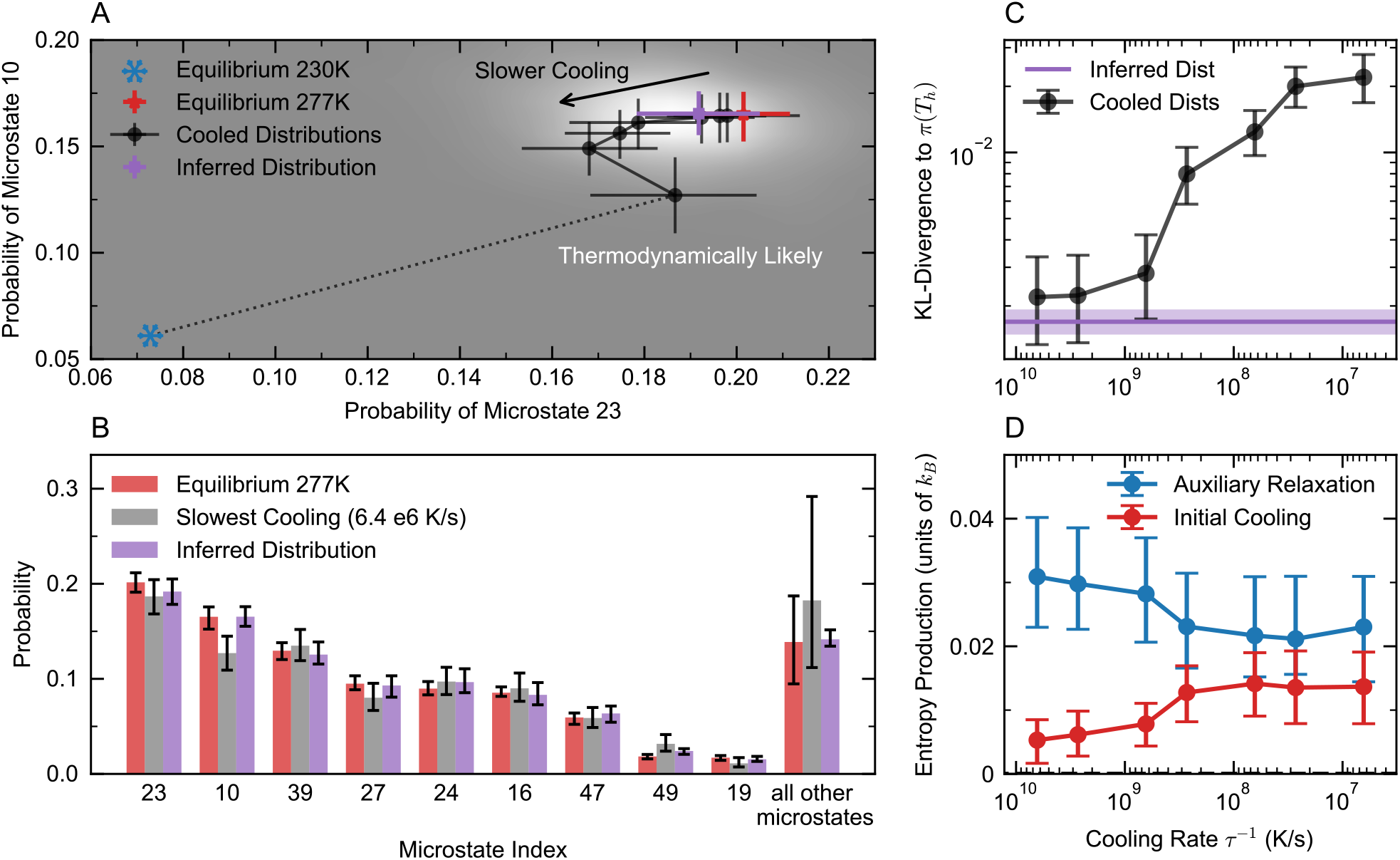
Using thermodynamic constraints to recover the equilibrium ensemble from cooled distributions. **A.** Projection of hot and cold equilibrium and cooled distributions onto the space of probabilities of microstates 10 and 23. **B**. Histogram showing the true equilibrium distribution, cooled distribution with the slowest cooling time, and inferred distribution using all cooling times in the calculation. **C**. KL-divergence for each nonequilibrium cooled distribution from the hot equilibrium distribution (black), as well as between the hot equilibrium distribution and the inferred distribution using all cooling times (purple). Light purple band indicates error bars. **D**. Entropy production for the initial cooling process (upper bound, red) and auxiliary relaxation process (blue) as functions of the cooling rate *τ*^−1^, computed using the inferred state probabilities and entropies.

Lighter white indicates a higher density of sampled thermodynamically-allowed distributions. We then obtained an estimated equilibrium distribution by taking the 10% of the thermodynamically allowed region closest to the fastest-cooled distributions, which provided the purple estimates with uncertainty in Fig. 6A and B showing the inferred distribution.

To demonstrate the utility and validity of this thermodynamic recovery method, we first analysed a toy model: overdamped diffusion of a Brownian particle in a two-dimensional potential energy landscape with six local minima (Fig. S11A). For this model system, which is sufficiently complex to show non-monotonic evolution of certain state probabilities with cooling time (Fig. S11B), our method accurately recovered the full equilibrium ensemble using only the collection of simulated cooled ensembles (Fig. S11E), obtaining a better estimate for the equilibrium ensemble than even the fastest simulated cooling time (Fig. S11F).

We then applied the thermodynamic recovery method to our simulations of the Trp-cage protein to infer equilibrium state populations. Fig. 6A illustrates the thermodynamic recovery method applied to the microstate MSM, showing a two-dimensional projection of the thermodynamically allowed space of equilibrium ensembles, which contains the true equilibrium ensemble. This visualization shows the distribution moving away from the thermodynamically allowed region as the cooling time increases. The thermodynamic recovery method produced an estimate of the distribution for the initial hot equilibrium ensemble from the trajectory of cooled distributions. We recovered the true equilibrium probabilities for the nine most probable microstates in the MSM within uncertainty (Fig. 6B), and found that the inferred distribution is closer to the true equilibrium, as quantified by KL-divergence, than even the fastest-cooled distributions (Fig. 6C). The inferred entropy productions for both the initial cooling and auxiliary relaxation processes are indeed non-negative, and monotonic with cooling time as required by our thermodynamic constraints (Fig. 6D).

Since our fastest cooling rates are faster than realistic cooling processes used in cryo-EM, we also tested the recovery process using only data from the three slowest cooling runs. As in the case of using all coolings, our recovery method successfully recovered an inferred distribution closer to the true equilibrium than any of the three slowest-cooled distributions (Fig. S12). While our thermodynamic recovery method performs best when multiple rank-ordered cooling times are available, it can still be applied using only data from a single cooling time by using only the non-negativity constraints on entropy production Eq. (2). We tested this approach using only the data from the critical cooling rate, and found that while uncertainties are large, the resulting inferred distribution is nonetheless closer to the true hot equilibrium than the cooled distribution (Fig. S13). These results show that our method can be applied even when multiple cooling rates are not available.

## Discussion

Through extensive molecular simulation and a computational non-equilibrium statistical mechanics approach, this work creates a foundation for understanding the perturbative effects of the vitrification process in cryo-EM. First, we have shown that water glass formation is unperturbed by the presence of the Trp-cage, and likely other biomolecules as well. We found that susceptibility to annealing during cooling depends on a state’s kinetic connectivity, with kinetically isolated states remaining more robust than well-connected ones. Perhaps most importantly, our results show that even kinetically active conformations present at 277 K will be observed, though perhaps not with the correct populations, in the cooled ensemble. We build on this result to propose a thermodynamic inference approach to recover equilibrium populations from the cryo-EM ensemble. Thus we have shown that recovering the correct ensemble populations from cryo-EM, a task with implications ranging from drug design to understanding the basic thermodynamic drivers of key biological processes, is possible.

Despite these strong conclusions, this work does not come without caveats. As with all classical simulations, we are limited by general force field accuracy, and by design choices in biopolymer force fields; no protein force field we are aware of is tuned for accuracy in supercooled water. In the case of Trp-cage, our choice of amber03w for the protein forcefield was based on previous validation that it exhibited folding behaviour consistent with experiment, and also showed a cold-denatured state (Fig. 3) [34–36]. Though no water model exhibits an accurate temperature for the glass transition of water, TIP4P/ICE represented the best compromise between cost and accuracy at 277 K and in the supercooled liquid [25]. While TIP4P/ICE exhibits a higher apparent *T*_*g*_ than real water, the more relevant comparison here is the temperature at which dynamics enter the regime where larger conformational rearrangements become strongly hindered. Real water shows a dynamical crossover near 233 K, and TIP4P/ICE places the corresponding crossover near 245 K [25, 37]. Proteins likewise undergo a dynamical transition around 240 K, associated with suppression of hydration-water translational diffusion and large-amplitude anharmonic motions [38]. Taken together, this suggests that our choice of *T*_*c*_ captures a similar dynamical regime to real vitrification, even if the transition temperature is shifted upward in TIP4P/ICE.

By inspecting water mobility and bulk rate-dependent density-temperature equations of state with and without protein, we establish that the system is large enough that Trp-cage does not materially alter water vitrification at ambient pressure in our simulations. While this result is an important control for our study, it is also suggestive. Because our protein concentrations are high (approximately 17 mM) relative to those used in cryoEM, one might suppose that if perturbations to glass formation were likely from solutes like proteins, we would have seen them. Therefore, it is reasonable to suspect that other changes to the solvent composition, such as low to moderate ionic strength, buffers, or other macromolecules at experimental concentrations, will likewise not perturb water glass formation, in agreement with experimental predictions of the effect of solutes on water vitrification [39].

The sampling required to achieve our results, 50.4 ms of total simulation time for a 20-residue protein, highlights a key limitation of this computational approach. This sampling includes 5.4 ms of unbiased simulation at 277 K, 1.1 ms of TREX, and 44 ms for the non-equilibrium cooling simulations, of which 18.4 ms were of the slowest cooling rate. This does imply that studying real cryo-EM systems through direct simulation of the cooling, as we have done here, is likely unfeasible given current computing resources. The unbiased dataset resulted in a well-sampled counts matrix for our MSM, with the least connected state still sampling 132,455 out transitions and 134,714 in transitions in aggregate. Despite the extensive unbiased sampling from an already converged TREX starting distribution and using modern MSM approaches, we could not initially build an MSM that was thermodynamically consistent with our cooling simulations or the state probabilities from histogramming our TREX data. This motivated using the TREX to constrain our MSM which in turn led to a thermodynamically consistent MSM with our cooling data. These findings suggest that thermodynamic constraints may prove useful more generally as a tool to rule out implausible MSM candidates.

Our extensive MD simulations establish quantitative relationships between biomolecular kinetics and vitrification-induced perturbations in cryo-EM. States with event times exceeding the duration of cooling are well preserved during vitrification. At the experimental critical rate (6.4 × 10^6^ K/s), for our system, this corresponded to states with event times around 7 µs. While faster-exchanging states exhibit measurable population shifts, their extent is a function of the state event time and the cooling duration. These results imply that the choice of most cryo-EM practitioners to equilibrate their samples at 277 K before vitrification is a good one, because they are minimizing cooling artefacts by minimizing the cooling duration. Our results showing that for a fairly fast exchanging system, Trp-cage, states from 277 K are well-preserved even at the slowest cooling rates. However, for cryo-EM practitioners interested in the states of their system at higher temperatures, say 25 ^°^C or 298 K, the results presented here might not hold, as the cooling from 298 K to 277 K could result in pronounced changes in the ensemble populations.

Previous simulation work on vitrification [18] established the feasibility of studying vitrification with MD, though their large system size necessitated cooling rates 1000-fold faster than experiments, and the precise critical cooling rate had yet to be put on firm experimental ground. Other simulations studying the impact of cryogenic temperatures on actin have recently been reported; like our study, preferences for certain configurations are observed to be sensitive to temperature [40]. Our focus on Trp-cage enabled exhaustive sampling at experimentally realistic rates, revealing how conformational kinetics quantitatively determine preservation. Because our approach to estimating the sensitivity of a state to annealing is based on all the transition paths available to a molecule, it is hard to compare it to earlier estimates by Bock and Grubmüller, who find that barriers below ∼10 kJ/mol can be traversed during cooling [18], thus framing the problem in a quasi two-state fashion.

It has been previously shown that the rapid cooling used in most crystallography for structure determination is slow enough to permit rearrangements of the side chains of the crystallized protein away from the room temperature ensemble [41–43]. This cooling usually happens over *>*1 ms, and is affected by the different solvent conditions as well as the protein crystal itself. We also observed side chain distribution changes within Trp-cage due to cooling (Fig. 5A, Fig. S9). This suggests that the cooling duration in cryo-EM is long enough to perturb side chain rotamer equilibrium distributions. Internal residue contacts remained stable within our simulations but this analysis is limited by the size of Trp-cage (Fig. 5C). Studying larger proteins undergoing vitrification might be required to confirm how internal and external residues are affected by vitrification.

Our thermodynamic recovery framework offers a path toward quantitative ensemble determination even when vitrification perturbs populations. While our method can be applied with only data from a single cooling rate, it performed best with data from multiple rank-ordered cooling rates. This could in principle be determined by estimating sample thickness across cryo-EM grids [18, 44]. A key advantage of our inference approach, however, is that it does not require precise cooling rate measurements, only rank ordering. Happily, it also does not directly require any notion of state kinetics. Our derivation assumed that protein volume does not change with temperature in a state-dependent manner (supported by Fig. S7), and that microstate energies and entropies are temperature-independent, which may not always be the case for coarse-grained states. Future work could seek to relax these assumptions to further generalize our approach. Realizing the potential of this approach requires experimental determination of molecular state populations from cryo-EM data, which is still an active area of methodological development [4, 45]. The combination of emerging heterogeneity analysis methods with our thermodynamic recovery framework could enable routine quantitative ensemble reconstruction from vitrified samples.

This study confirms that cryo-EM holds promise as a powerful quantitative technique for molecular biophysics. One area in which our results have direct implications is for time-resolved cryo-EM experiments [17] where experiments are approaching the microsecond time scale of vitrification itself [46] and cooling artifacts become potentially significant. Our framework provides both a criterion for when artifacts matter, and a method to correct for them when they do. Additionally, our recovery method inherently returns estimates for state-dependent internal energy and entropy of the protein that include uncertainty. These in principle are sufficient to compute temperature-dependent free energy landscapes for a subset of discrete states. Altogether, this work provides a path forward as conformational heterogeneity methods evolve, time-resolved methods move toward faster time resolution [46], and the understanding of the perturbative effects of rapid cooling outlined here will become increasingly relevant.

## Methods

### Molecular Dynamics

We used the Amber03w protein force field [34], which was previously shown to accurately capture cold denaturation in Trp-cage [14]. We chose the TIP4P/ICE water model [25] for its accurate reproduction of water’s physical properties near the freezing point compared to three-site models like TIP3P and competing four-point models [26]. All simulations were performed using GROMACS 2024.1. We solvated the first conformer from NMR structure PDB 1L2Y [47] in a periodic box with 3240 water molecules and one neutralizing chloride ion, resulting in a box with approximate edge length of 4.7 nm. A Nosé-Hoover thermostat with coupling time of 0.2 ps and a Parrinello-Rahman barostat with compressibility of 4.5 × 10^−5^ bar^−1^ and coupling time of 1 ps were used for all simulations [48–51]. All simulations were carried out at atmospheric pressure.

To obtain reference populations for Trp-cage at our temperatures of interest, we performed TREX across 90 replicas spanning temperatures from 230 K to 520 K [52]. Each replica was simulated for 12 µs, for an aggregate sampling time of 1.08 ms. The first 2 µs were excluded from analysis as equilibration. The final 10 µs from each replica were used to compute equilibrium populations at each temperature.

We seeded unbiased MD simulations at 277 K from 750 cluster centres determined by RMSD-based C_τ_ clustering using a Euclidean metric and the *k*-hybrid approach provided by the enspara library on our 277 K TREX trajectory [53]. Each cluster centre was initialised with a random velocity and we then simulated no fewer than 20 simulations of each cluster centre. We used a provisional MSM to target additional sampling to the 20 most populated prototype states, except these were seeded 10 random seeds belonging to each cluster. We also targeted some additional long-lived states that were poorly kinetically resolved within a prototype MSM. We ran most trajectories for 200 ns but increased the duration of some to 500, 750, 1000, and 2000 ns, for an aggregate dataset totalling 5.4 ms.

For all Trp-cage cooling simulations, excluding those at our slowest cooling rate, the simulations were seeded from the 750 centres of the RMSD clustering, and cooled by adjusting the set temperature of the thermostat in 1 K steps of increasing duration linearly interspersed between 277 and 230 K. Some cooling rates had an additional set of cooling simulations seeded from prototype MSMs, similar to as described above. This resulted in cooling trajectories of number and duration recorded in Supplementary Table 1. We ensured at least 10 replicas for all states across our cooling rates, though most states were sampled many more times (Supplementary Table 1, Fig. S16). In order to maximise our sampling for our compute budget for our slowest cooling rate, we prioritised certain molecular states that were more variable under cooling and states that were predicted to be central nodes for probability to pass through (See SI for additional details).

### Markov State Model Construction

To develop features, we used PyEMMA to extract all sine and cosine transformed dihedrals from our equilibrium 277K sampling [54]. Dimensionality reduction was performed with time-lagged independent component analysis (TICA) from the deeptime package [28, 55–58], by retaining 90% of the kinetic variance in these features. We used Koopman-reweighting to account for the relatively short length of individual trajectories [27]. We built multiple featurizations with several lag times ranging from 1 ns to 40 ns.

We used *k*-means clustering as implemented in deeptime to create MSM states. To determine the number of clusters, we performed a VAMP scan [8, 59, 60] to choose a “fittable” number of clusters. Briefly, we trained maximum likelihood MSMs on 50% train-test splits of featurized data, then rescored the test split with the VAMP decomposition of the trained model. We selected featurization hyperparameters to validate by picking the largest ones with concordant train and test VAMP2 scores (Fig. S15).

To validate our choice of model and state space, we constructed implied time scales plots using the Bayesian model estimator available in deeptime [29]. We assigned our TREX data to these clusters, then used the state probabilities at 277 K as a constraint on the stationary distribution during MSM fitting. We selected models with implied time scales showing smooth, monotonically increasing behaviour that reached plateau values as a function of lag time [61, 62]. Our final model has 50 microstates with a 25 ns TICA lag time and a 40 ns MSM lag time. More discussion of our modelling workflow can be found in the Supplement, in Section 3.

### Estimating the distribution of molecular states after nonequilibrium cooling simulations

To calculate the abundance of each state after cooling we counted the fractional abundance (or histogram weight) of each starting state based on its equilibrium probability toward the state that the trajectory ended in. This can be thought of as tracking packets of probability from beginning to end of the cooling process. For the microstates, this calculation was performed using

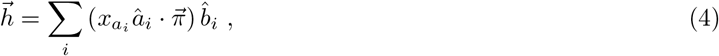

where *i* indexes the cooling runs, and 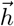 are the histogram weights for the microstates after cooling. The redundancy of seeding from each state is *x*_*a*_ = 1*/n*_*a*_ with *n*_*a*_ trajectories started from *a, â*_*i*_ and 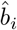 are indicators for start and end states of cooling run *i*, and 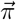 are the equilibrium populations.

We applied a Bayesian bootstrapping approach to obtain error estimates for these probabilities [63, 64]. To do this, we multiplied the above by a weight sampled from a Dirichlet distribution of the appropriate dimensionality as the prior, and produced 2000 (re)sampled cooling histograms, from which we reported means and 95% credibility intervals. For more details about the cooling simulations and seeding strategy, see SI.

### Categorizing change under cooling with hot equilibrium kinetics

To understand whether a state would exhibit change under cooling, we adapted a concept from the stochastic thermodynamics literature called the dynamical activity [21]:

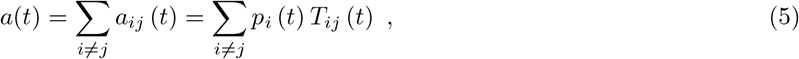

where *a*(*t*) is the dynamical activity at the model’s lag time *t*, with transition probabilities *T*_*ij*_(*t*), state prob-abilities *p*_*i*_(*t*), and the indices *i, j* correspond to the start and end states for the transition probabilities. At equilibrium and a particular lag-time τ, the contribution of state *k* to the activity will be

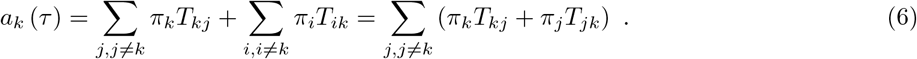

Because our MSM obeys detailed balance, the two summed terms are equal. We therefore scale the dynamical activity by 1/2. We further multiply by 0.05 to set a 5% threshold to define event time of state *k* at lag time τ as 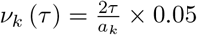.

This quantity is in units of time per unit probability, because the units of MSM step on the lag time and the dynamical activity both cancel. Thus, it is the time it would take, at equilibrium, for 5% of the total probability to flow out of state *k*.

To score the change in probability between equilibrium and cooled distributions on a per state basis, we computed the contribution of the state to the Kullback-Liebler divergence between the equilibrium distribution and the cooled distribution analysed

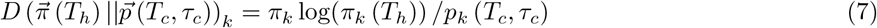

Where *T*_*h*_ is the hot temperature, *T*_*c*_ is the cold temperature, *π*_*k*_ is the hot equilibrium probability for state *k*, as above, and *p*_*k*_ (*T*_*c*_, τ_*c*_) is the probability of state *k* after nonequilibrium cooling over time τ_*c*_ to temperature *T*_*c*_.

### Thermodynamic constraints on cooling

We model the protein as a discrete-state stochastic system with *N* states labelled *{x*_*i*_ : *i* = 1, …,*N}*, subject to a time-dependent temperature *T* (*t*) varied according to a monotonically decreasing (but otherwise unknown) time-dependent protocol. The system is initialized at equilibrium at temperature *T* (0) = *T*_*h*_, and thus is initially described by the Boltzmann distribution

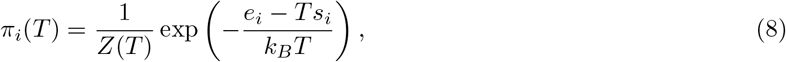

with *T* = *T*_*h*_. The distribution depends on the state-specific energies *{e*_*i*_*}* and entropies *{s*_*i*_*}*. We have implicitly assumed the protein volume to be independent of state, which is supported by analysis of the cooled protein ensemble (Fig. S7).

The temperature is then rapidly and monotonically decreased over a time τ to a lower temperature *T*_*c*_ *< T*_*h*_. At the end of this cooling process, the system is described by a nonequilibrium probability distribution *p*_*i*_(τ). We also consider an auxiliary relaxation process from *t* = τ to *t* = ∞, where the system relaxes to the equilibrium distribution *π*_*i*_(*T*_*c*_) at temperature *T*_*c*_.

To make progress, we assume that energies *{e*_*i*_*}* and entropies {*s*_*i*_} are temperature-independent, as is often done when modelling protein conformation distributions [65, 66]. This directly implies that the equilibrium distribution at the cold temperature *T*_*c*_ is given by the Boltzmann distribution, *π*_*i*_(*T*_*c*_).

We then assume that the system evolves according to a master equation with transition rates obeying local detailed balance at temperature *T* (*t*) for each time *t*. This allows us to apply the theory of stochastic thermodynamics [33] to quantify changes in energy and entropy as well as entropy production for both the initial cooling and auxiliary relaxation processes.

The ensemble-averaged energy *E* and entropy *S* of the system (here defined as only the protein) depend only on the probability distribution *p* over the states. They are given by

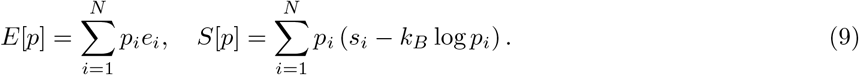

Here the energy is a weighted sum of state energies, while the entropy is a weighted sum of both state entropies and log-probabilities to account for coarse-graining [32]. We are interested in the entropy production due to the dynamics of the system. For a process from time *t*_*a*_ to *t*_*b*_, we can write

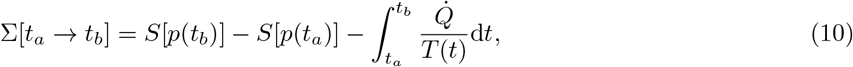

where 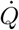 is the mean rate of heat flow into the system from its environment. Since no work is done during the cooling process, we have 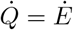, the mean rate of change of the system energy.

For the auxiliary relaxation process, since the temperature *T* = *T*_*c*_ is constant, the integral can be evaluated exactly to yield a simple expression for the entropy production

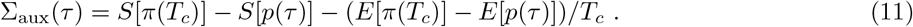

For the initial cooling process, since the temperature is time-dependent it is not generally possible to evaluate the integral, we proceed by assuming the mean energy decreases monotonically with temperature (such that *Ė /T* (*t*) ≥ *Ė /T*_*c*_, ∀ 0 ≤ *t* ≤ τ), which allows us to place an upper-bound (UB) on the entropy production

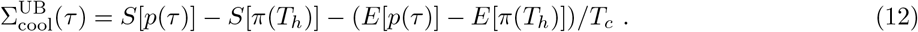

Conveniently, both entropy productions, requiring only differences in energies and entropies, do not depend on the (unknown) partition functions for the hot and cold equilibrium distributions.

Both the initial cooling and auxiliary relaxation processes must satisfy the second law of thermodynamics, Eq. (2). This provides two constraints for each cooling time τ. We then leverage having distributions at multiple cooling rates by comparing entropy production of the cooling and auxiliary processes given different cooling times. In particular, the longer the cooling time τ, the closer the cooled distribution *p*(τ) is to the cold equilibrium *π*_*c*_, and conversely the farther it is from *π*_*h*_. Thus, we conjecture (supported by numerical experiments, previous work studying nonequilibrium cooling [67–69], and a small-τ proof in the SI) that the entropy productions of the two processes are monotonic in τ. In particular, for two different cooling times τ_1_ *<* τ_2_, giving Eq. (3). This provides additional thermodynamic constraints, which become more useful as the density of cooling times increases. For a total of *N* cooling times, monotonicity gives 2(*N* − 1) additional thermodynamic constraints.

To obtain an estimate for the hot equilibrium distribution *π*(*T*_*h*_), we use all cooled distributions *{p*(τ)*}*, and employ a differential evolution algorithm to sample the region of thermodynamically likely values for the state probabilities *{π*_*i*_(*T*_*h*_)*}* and microstate entropies *{s*_*i*_*}*. We then determine the 10% of the thermodynamically allowed region closest to the fastest available cooling times, and use the center of mass of the resulting samples along with standard deviations as our estimates of each state probability. Our code for performing this analysis is available on Github.

## Acknowledgements

The Flatiron Institute is a division of the Simons Foundation. This project benefitted greatly from the technical support of the Scientific Computing Core of the Flatiron Institute and the expertise of Geraud Krawezik in particular. A portion of the computations presented in this study were performed on Folding@Home and we thank the participants of this platform for their contributions. The authors thank R. Justin Lindsay for assistance with performing the replica exchange simulations in this study as well as Sukrit Singh and Erik Thiede for feedback on the manuscript. LGS thanks Greg Bowman and Blanton Tolbert for support during this study, and Alan Grossfield for helpful conversations. MPL thanks Steven Blaber (UBC Physics) for helpful discussions, and Yale and NSERC for fellowship support. SK acknowledges funding from EPSRC (EPSRC grant numbers: EP/X035603, EP/V030779 (SK)) and SK acknowledges St Anne’s College. RC acknowledges funding from MRC (MRC grant numbers: MR/N013468 and MR/W00673), Magdalen College and Oxford Department of Biochemistry. Some authors made use of Claude Code and ChatGPT/Codex for organizing analysis pipelines, and for refining plotting scripts. All such work was manually reviewed for correctness.

## Notes

### Competing Interest Statement

The authors have declared no competing interest.

