## Supplementary Information for "Cooling fast and slow: Characterising the effects of vitrification in cryo-EM and the subsequent recovery of equilibrium populations"

#### 1 Glass Transition Temperature Determination

The glass transition temperature ($T_g$) was determined from MD cooling simulations using a hyperbola-based regression method adapted from Patrone et al.[30] This approach provides an objective, reproducible estimate of $T_g$ that avoids the subjective identification of asymptotic temperature regimes inherent in traditional bilinear fitting methods. We performed isobaric (NPT) cooling simulations at three cooling rates spanning three orders of magnitude: $6.4 \times 10^9$, $6.4 \times 10^8$, and $6.4 \times 10^7$ K/s. Systems were cooled linearly from 277 K to 80 K, below the experimental cryo-EM imaging temperature. We calculate ($T_g$) for two system types: pure TIP4P/ICE water and our Trp-cage protein system. At each temperature during cooling, the enthalpy was computed by averaging over 1 K windows.

Rather than fitting linear functions to subjectively identified high- and low-temperature regimes, we fit all enthalpy-temperature data simultaneously to a smooth hyperbola function:

$$H(T) = H_0 + a(T - T_0) + b\sqrt{(T - T_0)^2 + e^c} \tag{13}$$

where $H_0$ is the enthalpy offset, $T_0$ is the temperature at the transition centre, $a$ is the average slope across the transition, $b$ controls the difference between asymptotic slopes, and $c$ determines the sharpness of the transition. As $T \to -\infty$, this function approaches the glassy regime with slope $(a - b)$, while as $T \to +\infty$, it approaches the liquid regime with slope $(a + b)$. The hyperbola centre $T_0$ serves as the estimate of $T_g$. The five parameters $(H_0, T_0, a, b, c)$ were determined by non-linear least-squares fitting.

To objectively identify the glass transition region, we computed the derivative of the hyperbola:

$$\frac{dH}{dT} = a + \frac{b(T - T_0)}{\sqrt{(T - T_0)^2 + e^c}}. \tag{14}$$

The transition region was defined as the temperature range over which the derivative has converged to less than 90% of its asymptotic values. Specifically, we defined the percent convergence function:

$$P(T) = \frac{1}{2} + \frac{1}{2}\frac{(T - T_0)}{\sqrt{(T - T_0)^2 + e^c}} \tag{15}$$

and identified the transition region bounds $T_-$ and $T_+$ as the temperatures where $P(T) = 0.1$ and $P(T) = 0.9$, respectively. This criterion ensures that the simulated data samples both glassy and liquid regimes, validating the applicability of the hyperbola model.

Within-simulation uncertainty was estimated using bootstrap resampling. For each cooling rate, we performed 1000 bootstrap iterations. In each iteration, synthetic enthalpy data were generated by adding Gaussian noise to the fitted hyperbola:

$$H_{\text{synthetic},\ j} = H(T_j;\ \hat{\theta}) + \epsilon_j \tag{16}$$

where $\hat{\theta} = (H_0, T_0, a, b, c)$ are the fitted parameters, and $\epsilon_j \sim \mathcal{N}(0, \sigma_j^2)$. The hyperbola was refit to each synthetic dataset, yielding a distribution of $T_g$ estimates. The standard deviation of this distribution provides the within-simulation uncertainty.

##### 1.0.1 Representative Glass Transition Calculation

Figure S2 illustrates the hyperbola regression analysis for a representative cooling simulation at $6.4 \times 10^7$ K/s. Figure S2A shows the enthalpy-temperature data with the fitted hyperbola (black dashed line), the identified $T_g$ (red vertical line), and the transition region (shaded area). The hyperbola smoothly interpolates between the glassy and liquid regimes, providing an objective fit that uses all available data points rather than relying on subjective regime selection.

Figure S2B displays the derivative $dH/dT$ as a function of temperature, clearly showing the sigmoidal transition between the two asymptotic slopes (green and pink dashed lines). The rapid change in slope occurs within the identified transition region, confirming that $T_g$ corresponds to the inflection point of the enthalpy curve. The width of the transition region ($T_+ - T_-$) provides a measure of the breadth of the glass transition. While classical water models like TIP4P/ICE do not simultaneously reproduce both room-temperature and cryogenic properties of water with high fidelity, the glass transition behaviour is clearly observed and well-characterized by the hyperbola method.

Notably, the identified transition region begins near 230 K (specifically, $T_+ = 230.6$ K for this cooling rate), which provides independent validation for our decision to terminate the extensive protein cooling simulations at this temperature. The hyperbola analysis confirms that 230 K marks the onset of glassy dynamics in TIP4P/ICE water at these cooling rates, where the solvent begins to deviate from liquid-like behaviour and transitions toward the glassy state. This is consistent with our observations that water self-diffusion becomes extremely low below 230 K and protein state populations change minimally beyond this temperature (Fig. S3). The convergence of these independent lines of evidence, the thermodynamic (via the transition region), dynamical (via diffusion), and kinetic (via protein state populations), strongly supports 230 K as a rational stopping point that captures the essential physics of vitrification while substantially reducing computational cost.

Figure S2C shows the bootstrap distribution of $T_g$ estimates (histogram) overlaid with a fitted normal distribution (blue line). The narrow distribution confirms that the $T_g$ estimate is robust with respect to statistical fluctuations in the enthalpy data.

This analysis was repeated for three cooling rates ($6.4 \times 10^9$, $6.4 \times 10^8$, and $6.4 \times 10^7$ K/s) and for both pure water and Trp-cage-containing systems. As expected from equilibrium thermodynamics, $T_g$ decreased monotonically with decreasing cooling rate. Critically, we found that $T_g$ values for the protein-containing systems were statistically indistinguishable from those of pure water at each cooling rate (main text Fig. 4 inset), demonstrating that a single small protein molecule does not measurably perturb the glass transition of the surrounding solvent. This corroborates the assumption that the bulk water vitrification proceeds essentially independently of the protein dynamics during rapid cooling.

The hyperbola method provides several advantages over traditional bilinear fitting: (1) it eliminates subjective decisions about which data points belong to asymptotic regimes, (2) it uses all available data simultaneously rather than discarding transition-region points, (3) it provides a smooth, differentiable model suitable for uncertainty propagation, and (4) it includes built-in criteria for assessing whether the data adequately samples both regimes. These features make the method particularly well-suited for automated, reproducible analysis of simulation data across different cooling rates and system compositions.

#### 1.1 Cooling rates and trajectory information

**Table 1** Summary of nonequilibrium cooling simulations.

| Cooling rate | Simulation time between 277 K and 230 K (ns) | number of trajectories | aggregate sampling (ms) |
|---|---|---|---|
| $6.4 \times 10^9$ K/s | 7.3 | 7550 | 0.05 |
| $2.7 \times 10^9$ K/s | 17.9 | 9500 | 0.17 |
| $6.4 \times 10^8$ K/s | 73.4 | 7550 | 0.55 |
| $2.7 \times 10^8$ K/s | 179.4 | 7550 | 1.35 |
| $6.4 \times 10^7$ K/s | 734.4 | 13539 | 9.94 |
| $2.7 \times 10^7$ K/s | 1793.9 | 7550 | 13.54 |
| $6.4 \times 10^6$ K/s | 7343.8 | 2500 | 18.36 |

**Table 2** Pilot simulations to 196 K (not analysed in main text)

| Cooling rate | Simulation time between 277 K and 196 K (ns) | number of trajectories | aggregate sampling (ms) |
|---|---|---|---|
| $6.4 \times 10^7$ K/s | 1250 | 5341 | 6.68 |

### 2 Protein Volume Analysis

#### 2.1 Methods

Protein volumes were calculated from MD trajectories using GROMACS 2024.1 [CITE GROMACS]. For each trajectory, the molecular volume was computed using the `gmx sasa` tool with a probe radius of 0.0 Å, which calculates the volume enclosed by the van der Waals surface of the protein atoms. This approach provides a direct measure of the excluded volume occupied by the protein molecule. Volume calculations were validated against independent Voronoi tessellation analysis using van der Waals radii, which gave consistent results.

The analysis was performed for all distinct protein conformational clusters (microstates) as identified through our MSM, each sampled with 10 independent replica simulations, at three different cooling rates. For each cluster, protein volumes were calculated at temperature plateaus spaced at 1 K intervals throughout the cooling protocol (276 K to 80 K for $q_c = 6.4 \times 10^9$ and $q_c = 6.4 \times 10^8$ K/s and 276 K to 230 K for $q_c = 6.4 \times 10^7$ K/s). The 10 replica simulations within each cluster were averaged to obtain cluster-resolved volume estimates at each temperature. To quantify cooling-induced volume changes while accounting for the structural diversity across clusters, we employed a normalized analysis. For each cluster, the protein volume at each temperature was normalized relative to its high-temperature reference value:

$$\Delta V_{\mathrm{rel}}(T) = \frac{V(T) - V_{\mathrm{ref}}}{V_{\mathrm{ref}}} \times 100\% \tag{17}$$

where $V_{\mathrm{ref}}$ is the cluster-specific reference volume at $T = 277$ K and $V(T)$ is the volume at temperature $T$. This normalization removes baseline volume differences between clusters arising from their different structural characteristics, allowing direct comparison of cooling-induced volume changes across all microstates.

#### 2.2 Analysis

Figure S7 shows the distribution of normalized volume changes for all three cooling rates. The left panels display violin plots showing the probability density of volume changes across all microstates, while the right panels show individual cluster trajectories coloured by their final volume change. For the faster cooling rates, two distributions are shown. The full temperature range (276 K $\rightarrow$ 80 K, lighter colour) and a restricted range (276 K $\rightarrow$ 230 K, darker colour) to enable direct comparison with the slower cooling rate ($6.4 \times 10^7$ K/s).

This reveals that protein volume remains remarkably constant across the entire temperature range studied. The distributions of relative volume changes are centred near zero for all cooling rates, with median changes of $-0.09\%$ ($q_c = 6.4$ K/s), $-0.01\%$ ($q_c = 0.64$ K/s), and $-0.02\%$ ($q_c = 0.064$ K/s) over the 276 K $\rightarrow$ 230 K range. Even for the full cooling range to 80 K, the median changes remain small: $-0.15\%$ ($q_c = 6.4$ K/s) and $-0.06\%$ ($q_c = 0.64$ K/s). The distributions show characteristic widths with extremal values extending to approximately $\pm 1.25\%$, demonstrating that even the most structurally responsive microstates exhibit only modest volume changes during cooling. The violin plot representation illustrates that the majority of configurations experience minimal volume change.

The right panels of Figure S7 display individual cluster trajectories, revealing heterogeneous volume responses across different microstates. Trajectories are coloured according to their final volume change using a diverging red-blue colour map (red indicates expansion, blue indicates contraction). While the ensemble-averaged response is minimal, individual clusters show systematic trends, with some microstates undergoing slight contraction (blue trajectories) and others showing modest expansion (red trajectories). The top three clusters with the largest magnitude changes (highlighted with thicker lines) demonstrate that even the most responsive microstates change by less than 1.5%.

### 3 Markov State Model construction and sampling

Here we would like to elaborate on the remarks we made in regard to the construction of our Markov State Model (MSM) in the main text. Making MSMs is challenging despite advances in data-driven approaches to choosing hyperparameters. We wanted to make this MSM in a somewhat conventional way, where our choice of states was derived from kinetically informed features and validated using kinetic heuristics for quality; we also found we needed to constrain its stationary distribution to the histogrammed distribution for the same states from our Temperature replica exchange (TREX) in order to get a model that was consistent across our data and seemed to behave well in the context of our recovery approach.

#### 3.1 Sampling

To improve our sampling of the equilibrium ensemble at 277 K we used a very fine geometric clustering based on C-alpha RMSDs to partition the TREX data into states. We used frames corresponding to this partitioning of the temperature replica exchange dataset to initialize trajectories for these simulations. The objective in doing this was to make sure we had overlapping and detailed coverage of the full conformational diversity spanned by our TREX. We needed to get this coverage from contiguous and unbiased trajectories in order to make a kinetic model based on counting transitions (in our case an MSM).

To initialize a simulation, we picked a sample frame that was assigned to a given cluster index, then we randomized the velocities for that configuration and ran the simulation under the specified conditions. By running replicas starting from these states we were able to generate a large collection of trajectories that, when clustered in a kinetically weighted feature space, transited many states, thus giving us many opportunities to estimate the probability of transition between any pair of states. This approach to sampling fit well with our computational resource, which was the `PREEMPT` queue at the Flatiron Institute's main cluster, 'rusty'. We were able to utilize CPU resources that would have otherwise been idle, and we were able to avoid getting booted off of nodes very often by running many short trajectories that did not have a long wall-time, resulting in the swarm-of-trajectories dataset shape that is the classic application of MSMs to molecular dynamics modeling.

This was the approach taken both for extending sampling at 277 K and also for simulating cooling. In the case of simulated rapid cooling we turned the temperature down on the thermostat in a linear ramp with one kelvin increments between 277 K and 230 K, our chosen end temperature. Changing the number of simulation steps taken before reducing the temperature by a degree effectuated the various cooling rates we sought to simulate. We found that changing the temperature continuously per simulation step was not feasible, as the thermostat couldn't keep up with the system as it cooled. This resulted in a temperature that fluctuated significantly more than expected within a short window of the trajectory.

#### 3.2 Feature and Model Selection

To cluster this dataset using kinetically relevant features, we turned to time independent component analysis (TICA) followed by KMeans clustering on the sine and cosine transformed rotatable dihedrals for every residue. This permitted a data-driven dimension reduction in the number of features; we chose to only include the largest TICA modes that captured 90% of the kinetic variance in the features. After both of these steps, we used the VAMP-2 score with 50% train-test splits on our feature trajectories to determine a fit-able number of states for KMeans (see Fig. S15).

In model selection we noted a one rare (in the TREX data it was only 'found' in one trajectory and was a teeny fraction of the 277 K ensemble) state that, when used as a seed, produced no out-transitions in the counts matrix. To investigate this further, we vastly extended our sampling of this state; it was a curious looking Arginine salt-bridge that held most of the protein in a rigid conformation. After extensive focused seeding from this state, totalling hundreds of µs, we eventually saw an out transition, but no in-transitions. We also noted, however, that instead of trimming this state our kinetic clustering was prioritizing it, and attempting to estimate its probability. Because ergodic trimming ought to have eliminated the state centred around this conformation, and is common practice in MSM construction, we decided to manually exclude trajectories containing this state from our model construction [70]. This did not change our conclusions about how the event time can categorize cooling, but changed the slowest timescale we observed (because the transitions were estimated into and out of this state as being very infrequent). We note that to our surprise this state was not attributed high kinetic variance, and its inclusion didn't add uncertainty in the longest implied timescales. As always, when making MSMs, extreme caution is advised.

##### 3.3 MSM construction

As our TREX data provided an estimate of the equilibrium distribution at 277 K, we used this to constrain the stationary distribution of the MSM. The TREX trajectory was projected into the same kinetic feature and state space, and uncertainty in the stationary distribution was estimated by block bootstrapping the final 10 µs of sampling to ensure sufficient equilibration. We then propagated this uncertainty into our MSM by estimating a separate reversible Bayesian MSM for each of 50 bootstrap samples of our TREX-derived stationary distribution. From each constrained Bayesian MSM, we drew 100 posterior samples, giving 5000 total samples which formed our posterior ensemble.

#### 4 Seeding of cooling simulations at $6.4 \times 10^6$ K/s

We first generated an initial round of $6.4 \times 10^6$ K/s cooling simulations, with between 3 and 20 simulations per MSM state. Based on our MSM, we computed a PageRank-style stationary distribution on the transition matrix to define a target distribution of starting states that reflected both stationary weight and network connectivity [71]. We then quantified how under- or over-sampled each state was relative to this target. We combined this with a simple metric that used the cooling data itself, that penalised poorly covered states and those with heterogeneous outgoing transition patterns in the pre-cooled ensemble. This metric worked by ensuring there were at least 10 of every state, and that states with high Shannon entropy in outcome were given more weight. These scores were rescaled and combined into a single priority value per state, and our fixed simulation budget was allocated in a single "one-shot" step in proportion to these priorities, using integer rounding to preserve the total budget exactly.

#### 5 Details of Recovering the Equilibrium Distribution from Quenched Ensembles.

##### 5.1 Detailed Derivation of Thermodynamic Constraints

Here we provide an expanded version of the derivation of thermodynamic constraints in the main text.

We model the protein as a discrete-state stochastic system with $N$ states labelled $\{x_i : i = 1, ..., N\}$, subject to a time-dependent temperature $T(t)$ varied according to a monotonically decreasing (but otherwise unknown) time-dependent protocol. We assume that the system evolves according to a master equation with transition rates ($k_{ij}$ denotes the transition rate from state $j$ to state $i$) obeying local detailed balance at temperature $T(t)$ for each time $t$:

$$\frac{k_{ij}(t)}{k_{ji}(t)} = \exp\left(-\frac{e_i - e_j}{T(t)}\right). \tag{18}$$

The dynamics of the probability distribution are then described by the master equation

$$\frac{\mathrm{d}}{\mathrm{d}t}\boldsymbol{p}(t) = \boldsymbol{R}(t)\boldsymbol{p}(t), \tag{19}$$

where $\boldsymbol{R}(t)$ is the time-dependent transition rate matrix with entries given by

$$R_{ii}(t) = -\sum_{j\neq i} k_{ji}(t), \tag{20a}$$

$$R_{ij}(t) = k_{ij}(t). \tag{20b}$$

We make no further assumptions about the transition rates.

The system is initialized at equilibrium at temperature $T(0) = T_h$, and because of local detailed balance is thus initially described by the Boltzmann distribution

$$\pi_i(T_h) = \frac{1}{Z_h} \exp\left(-\frac{e_i - T_h s_i}{k_B T_h}\right) . \tag{21}$$

The distribution depends on the state-specific energies $\{e_i\}$ and entropies $\{s_i\}$. In this derivation entropies have units of Boltzmann's constant $k_B$. We have implicitly assumed the protein volume to be independent of state, which is supported by analysis of the cooled protein ensemble (Fig. S7).

The temperature is then rapidly cooled over a time $\tau$ to a lower temperature $T_c < T_h$. We assume the temperature decreases monotonically. At the end of this rapid cooling process, the system is described by a nonequilibrium probability distribution $p_i(\tau)$. It will prove helpful to also consider an auxiliary relaxation process from $t = \tau$ to $t = \infty$, where the system relaxes to the equilibrium distribution $\pi_i(T_c)$ at temperature $T_c$.

To make progress, we assume that energies $\{e_i\}$ and entropies $\{s_i\}$ are temperature-independent, as is often done when modelling protein conformation distributions [65, 66]. This directly implies that the equilibrium distribution at the cold temperature $T_c$ is given by the Boltzmann distribution,

$$\pi_i(T_c) = \frac{1}{Z_c} \exp\left(-\frac{e_i - T_c s_i}{k_B T_c}\right) , \tag{22}$$

and is thus implicitly related to the hot equilibrium distribution.

The ensemble-averaged energy $E$ and entropy $S$ of the system (here defined as only the protein) depend only on the probability distribution $p$ over the states. They are given by [32]

$$E[p] = \sum_{i=1}^{N} p_i e_i \ , \tag{23a}$$

$$S[p] = \sum_{i=1}^{N} p_i \left(s_i - k_B \log p_i\right) \ . \tag{23b}$$

We are interested in the entropy production due to the dynamics of the system. For a process from time $t_a$ to $t_b$, we can write

$$\Sigma[t_a \to t_b] = S[p(t_b)] - S[p(t_a)] - \int_{t_a}^{t_b} \frac{\dot{Q}}{T(t)} \mathrm{d}t, \tag{24}$$

where $\dot{Q}$ is the mean rate of heat flow into the system from its environment. Since no work is done during the cooling process, we have $\dot{Q} = \dot{E}$, the mean rate of change of the system energy.

For the auxiliary relaxation process, since the temperature $T = T_c$ is constant, the integral can be evaluated exactly to yield a simple expression for the entropy production

$$\Sigma_{\mathrm{aux}}(\tau) = S[\pi(T_c)] - S[p(\tau)] - (E[\pi(T_c)] - E[p(\tau)])/T_c \ . \tag{25a}$$

For the initial cooling process, since the temperature is time-dependent it is not generally possible to evaluate the integral, we proceed by assuming the mean energy decreases monotonically with temperature (such that $\dot{E}/T(t) \geq \dot{E}/T_c$, for all $t$ such that $0 \leq t \leq \tau$), which allows us to place an upper-bound (UB) on the entropy production

$$\Sigma_{\mathrm{cool}}^{\mathrm{UB}}(\tau) = S[p(\tau)] - S[\pi(T_h)] - (E[p(\tau)] - E[\pi(T_h)])/T_c \ . \tag{26}$$

Conveniently, both entropy productions, requiring only differences in energies and entropies, do not depend on the (unknown) partition functions for the hot and cold equilibrium distributions.

Both the initial cooling and auxiliary relaxation processes must satisfy the second law of thermodynamics, i.e. $\Sigma_{\mathrm{cool}} \geq 0$ (and thus necessarily $\Sigma_{\mathrm{cool}}^{\mathrm{UB}} > 0$) and $\Sigma_{\mathrm{aux}} \geq 0$. This gives us two constraints for each cooling time $\tau$, for a total of $2N$ constraints given $N$ cooling times.

We can leverage having distributions at multiple cooling rates by comparing entropy production of the cooling and auxiliary processes for all of them. In particular, the longer the cooling time $\tau$, the closer the cooled distribution $p(\tau)$ is to the cold equilibrium $\pi_c$, and conversely the farther it is from $\pi_h$. Thus, we conjecture (supported by numerical experiments, and previous work studying nonequilibrium cooling [67–69]) that the entropy productions of the two processes are monotonic in $\tau$. In particular, for two different cooling times $\tau_1 < \tau_2$, we conjecture that

$$\Sigma[0 \to \tau_1] \leq \Sigma[0 \to \tau_2], \tag{27a}$$

$$\Sigma[\tau_1 \to \infty] \geq \Sigma[\tau_2 \to \infty]. \tag{27b}$$

In the next subsection we will prove these inequalities in the small-$\tau$ limit. These two inequalities provide additional thermodynamic constraints, and become more useful as the density of cooling times increases. For a total of $N$ cooling times, monotonicity gives $2(N-1)$ additional thermodynamic constraints. The inference problem is then reduced to finding the set of state energies $\{e_i\}$ and entropies $\{s_i\}$ that satisfy all thermodynamic constraints, given measurements of nonequilibrium cooled distributions $p(\tau)$ for several different cooling times $\tau$.

#### 5.2 Monotonic Entropy Production for Rapid Cooling

Here we prove that in the small-$\tau$ (rapid cooling) limit, the auxiliary entropy production $\Sigma_{\text{aux}}(\tau)$ decreases monotonically with the cooling time $\tau$, while the upper bound on the cooling entropy production $\Sigma_{\text{cool}}^{\text{UB}}(\tau)$ increases monotonically with $\tau$.

To compare different cooling times $\tau$, we assume that the corresponding temperature protocols are related by a time-rescaling of a single monotonic protocol shape. More specifically, we define $u \equiv t/\tau$, and assume that for all $\tau$, the cooling protocol satisfies

$$T(t) = T(u\tau) = \mathcal{T}(u), \tag{28}$$

where $\mathcal{T}$ is a monotonically decreasing function of $u \geq 0$ with $\mathcal{T}(0) = T_h$ and $\mathcal{T}(u \geq 1) = T_c$.

We now assume that $\boldsymbol{R}$ is only time-dependent through $T(t)$. This satisfies the assumption of generalized detailed balance, and is consistent with previous work on rapid nonequilibrium cooling in Cryo-EM [18]. We then write $\boldsymbol{R}(u) \equiv \boldsymbol{R}(T(t))$, so that the master equation becomes

$$\frac{\mathrm{d}}{\mathrm{d}u}\boldsymbol{p}_\tau(u) = \tau \boldsymbol{R}(u)\boldsymbol{p}_\tau(u), \tag{29}$$

with $\boldsymbol{p}_\tau(0) = \boldsymbol{\pi}(T_h)$. Here $\boldsymbol{p}_\tau(u)$ denotes the probability distribution at protocol time $u$ for a cooling process of duration $\tau$.

We next integrate the master equation and expand around $\tau = 0$ to obtain

$$\boldsymbol{p}_\tau(1) = \boldsymbol{\pi}(T_h) + \tau \underbrace{\int_0^1 \boldsymbol{R}(u)\boldsymbol{\pi}(T_h)\,\mathrm{d}u}_{\boldsymbol{A}} + \mathcal{O}(\tau^2). \tag{30}$$

Here we have defined $\boldsymbol{A}$ as the first-order perturbation to $\boldsymbol{\pi}(T_h)$, which must satisfy $\sum_i A_i = 0$ to ensure normalization of $\boldsymbol{p}(\tau)$.

We now consider the entropy production of the auxiliary relaxation process, which can be written as

$$\Sigma_{\text{aux}}(\tau) = k_B \sum_{i=1}^{N} p_i(\tau) \ln \frac{p_i(\tau)}{\pi_i(T_c)}. \tag{31}$$

Expanding around $\tau = 0$ using Eq. (30) gives

$$\Sigma_{\text{aux}}(\tau) = k_B D_{\text{KL}}[\boldsymbol{\pi}(T_h)||\boldsymbol{\pi}(T_c)] + k_B \tau \sum_{i=1}^{N} A_i \ln \frac{\pi_i(T_h)}{\pi_i(T_c)} + \mathcal{O}(\tau^2). \tag{32}$$

Since the first term is independent of $\tau$, we need only examine the first order correction.

We will approach this first-order correction term by considering a new auxiliary dynamics involving relaxation at a fixed temperature, which we will take to be $\mathcal{T}(u)$ for some $u \in (0,1]$. The master equation for such dynamics is

$$\frac{\mathrm{d}}{\mathrm{d}s}\boldsymbol{q}_u(s) = \boldsymbol{R}(u)\boldsymbol{q}_u(s), \tag{33}$$

where here $\boldsymbol{q}_u(s)$ is the probability distribution at time $s$ for a given choice of $u$. As initial conditions, consider $\boldsymbol{q}_u(0) = \boldsymbol{\pi}(T_h)$. Since $\boldsymbol{R}(u)$ obeys detailed balance at the fixed temperature $\mathcal{T}(u)$, the KL-divergence to $\boldsymbol{\pi}(\mathcal{T}(u))$ must be non-increasing,

$$\frac{\mathrm{d}}{\mathrm{d}s}D_{\mathrm{KL}}[\boldsymbol{q}_u(s)||\boldsymbol{\pi}(\mathcal{T}(u))] \leq 0 \tag{34}$$

for all $s \geq 0$. In particular this must be true at $s = 0$, so that

$$\sum_{i=1}^{N} [\boldsymbol{R}(u)\boldsymbol{\pi}(T_h)]_i \ln \frac{\pi_i(T_h)}{\pi_i(\mathcal{T}(u))} \leq 0. \tag{35}$$

Using the definition for the equilibrium Boltzmann distribution, we have

$$\ln \frac{\pi_i(T_h)}{\pi_i(\mathcal{T}(u))} = \frac{e_i}{k_B}\left(\frac{1}{\mathcal{T}(u)} - \frac{1}{T_h}\right) + \ln \frac{Z(\mathcal{T}(u))}{Z(T_h)}. \tag{36}$$

Since $\sum_i [\boldsymbol{R}(u)\boldsymbol{\pi}(T_h)]_i = 0$, this implies that for all $u \in (0,1]$ we have

$$\sum_{i=1}^{N} e_i [\boldsymbol{R}(u)\boldsymbol{\pi}(T_h)]_i \leq 0. \tag{37}$$

Now recall that the first order correction term for the auxiliary relaxation process entropy production is

$$\begin{aligned} k_B\tau \sum_{i=1}^{N} A_i \ln \frac{\pi_i(T_h)}{\pi_i(T_c)} &= k_B\tau \int_0^1 \mathrm{d}u \sum_{i=1}^{N} [\boldsymbol{R}(u)\boldsymbol{\pi}(T_h)]_i \ln \frac{\pi_i(T_h)}{\pi_i(T_c)} \\ &= \tau\left(\frac{1}{T_c} - \frac{1}{T_h}\right)\int_0^1 \mathrm{d}u \sum_{i=1}^{N} e_i [\boldsymbol{R}(u)\boldsymbol{\pi}(T_h)]_i\,. \end{aligned} \tag{38}$$

Since $T_c < T_h$, the term $\left(\frac{1}{T_c} - \frac{1}{T_h}\right) > 0$, and since Eq. (37) holds for all $u \in (0,1]$, the entire term is non-positive. This implies that so long as the second-order term in $\Sigma_{\mathrm{aux}}(\tau)$ can be neglected, we must have

$$\Sigma_{\mathrm{aux}}(\tau_1) \geq \Sigma_{\mathrm{aux}}(\tau_2), \quad \forall\, \tau_1 \leq \tau_2. \tag{39}$$

To show that $\Sigma^{\mathrm{UB}}_{\mathrm{cool}}$ is similarly monotonic with $\tau$, note that from the definitions of $\Sigma_{\mathrm{aux}}$ and $\Sigma^{\mathrm{UB}}_{\mathrm{cool}}$,

$$\Sigma_{\mathrm{aux}}(\tau) + \Sigma^{\mathrm{UB}}_{\mathrm{cool}}(\tau) = S[\pi(T_c)] - S[\pi(T_h)] - \frac{E[\pi(T_c)] - E[\pi(T_h)]}{T_c}, \tag{40}$$

which is independent of $\tau$. Therefore

$$\frac{\mathrm{d}}{\mathrm{d}\tau}\Sigma^{\mathrm{UB}}_{\mathrm{cool}}(\tau) = -\frac{\mathrm{d}}{\mathrm{d}\tau}\Sigma_{\mathrm{aux}}(\tau). \tag{41}$$

Thus since $\Sigma_{\mathrm{aux}}$ decreases monotonically with $\tau$, $\Sigma^{\mathrm{UB}}_{\mathrm{cool}}$ must similarly increase monotonically with $\tau$.

#### 5.3 Recovery Algorithm

Here we outline the implementation details of the thermodynamic recovery algorithm. To begin, we sampled points uniformly in the space of valid $N$-state probability distributions and state-dependent entropies. State probabilities are bounded on $[0,1]$ subject to the requirement of normalization, while, lacking strong priors for state entropies, we took the fairly permissive range $s_i \in [0, 5]$ (our results are not dependent on this choice). For each sampled point, we then iterated a differential evolution algorithm to evolve the state probabilities and entropies until all thermodynamic inequalities are satisfied. Specifically these constraints are main text equations 1 and 2, which together provide $2(2n-1)$ total constraints given $n$ distinct cooling times. We repeated this process for $10^4$ randomly-drawn initial points in order to sample the space of thermodynamically-allowed distributions. This provides the white region Fig. S11A, which is an exact boundary.

For the Trp-cage protein simulations error bars on the state probabilities after cooling were much larger, and thus the estimates of the thermodynamically likely region much noisier. Thus for display purposes we used KDE to fit the set of distributions determined to satisfy the thermodynamic constraints, which yields the white regions shown in Figs. 6A, S12A, S13A, and S14, which we denote "thermodynamically likely". To obtain an estimated equilibrium distribution we updated the fastest-cooled distributions towards the centroid of the thermodynamically allowed region by taking the 10% of the thermodynamically likely samples closest to the fastest-used cooled distributions, which yielded mean estimates for each probability with uncertainty. Our full code for implementing the recovery algorithm is available on github.

#### 5.4 Additional Results

As discussed in the main text, we initially validated the thermodynamic recovery method using a toy model: overdamped diffusion of a Brownian particle in a two-dimensional potential energy landscape with six local minima (Fig. S11A). We performed overdamped Brownian dynamics simulations of cooling from $T_h = 500$K to $T_c = 100$ K at 10 different cooling rates. Fig. S11B shows the probabilities of each state as functions of cooling time (in dimensionless simulation steps). As shown in Fig. S11C, the thermodynamics of the system obey both the second law ($\Sigma_{\text{cool}}(\tau) \geq 0$ and $\Sigma_{\text{aux}} \geq 0$), as well as our conjectured monotonicity constraints. Our method accurately recovered the full equilibrium ensemble (Fig. S11D and E), obtaining a better estimate for the equilibrium ensemble than even the fastest simulated cooling time (Fig. S11F).

While in main text Fig. 6 we showed the results of applying the thermodynamic recovery method to Trp-cage simulation results for all studied cooling times, we also tested the method using only the three slowest cooling times, which are more experimentally realistic. Figure S12 shows the results, namely that the inferred distribution is still closer to the true equilibrium than the fastest cooled distribution used. Figure S13 shows the same results using only the single slowest cooling.

Finally in Fig. S14 we show two-dimensional projections of the thermodynamic recovery for all pairs of states with probabilities greater than 1%, displayed both with and and without the cold equilibrium probabilities.

### 6 Supplementary figures

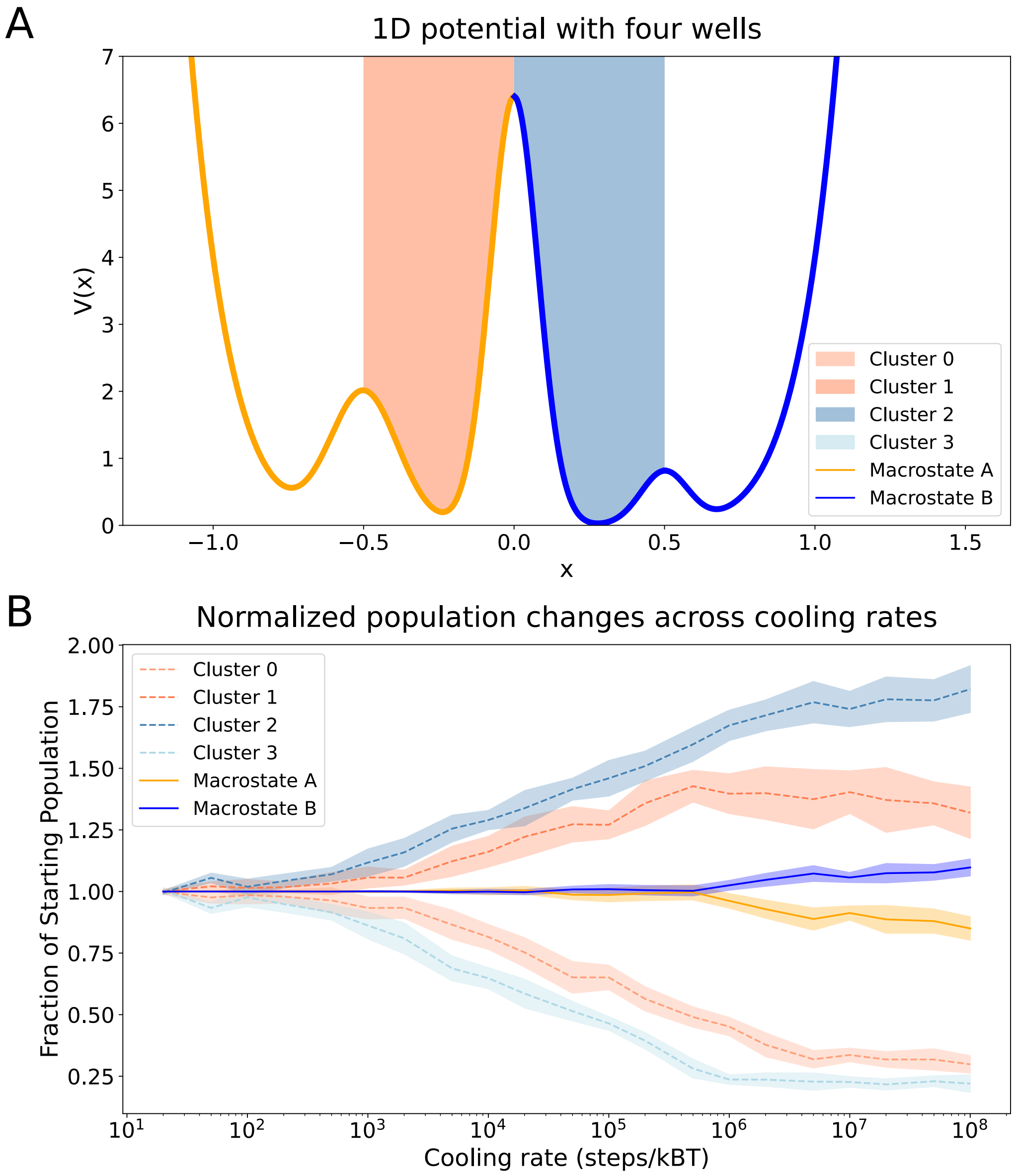


**Fig. S1 Non-equilibrium cooling simulations of overdampened Langevin dynamics on a one-dimensional four-well potential energy surface. A.** The potential energy surface is composed of four wells in one dimension. Two macrostates are separated by a large energy barrier. These macrostates are composed of two microstates each of which are separated by a smaller barrier. **B.** Using Langevin dynamics, we simulated the evolution of the system at different cooling rates down to 0 K. As a function of cooling rate, the quenched ensembles exhibit a non-monotonic behaviour as states are given more time to exchange with one another. The equilibrium distribution of this system at 0 K corresponds to 100% of the probability density being in the ground state (cluster 2), but this would require an infinitely slow cooling rate to achieve.

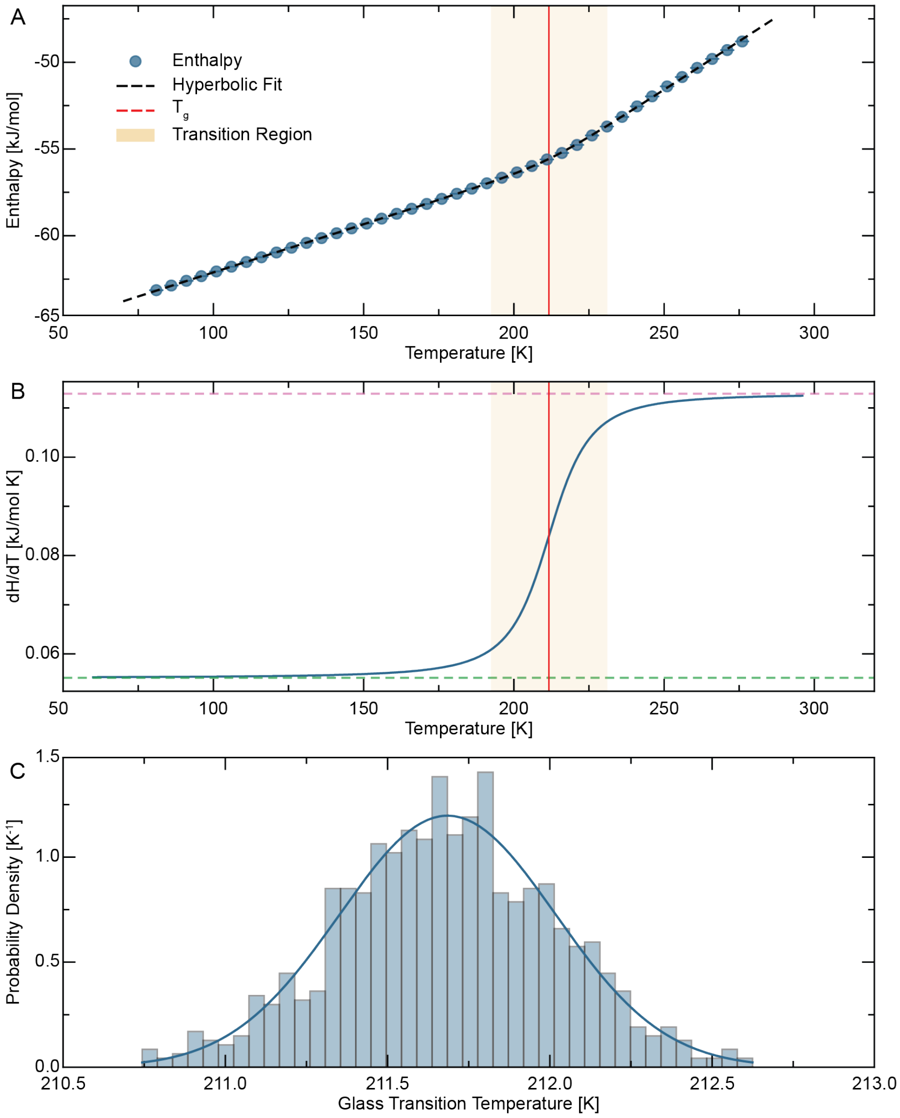


**Fig. S2 Hyperbola-based determination of glass transition temperature.** Representative analysis of determining $T_g$ using global hyperbola regression. **A.** Enthalpy per molecule as a function of temperature (blue circles, showing every fifth data point from the 1 K plateau-averaged values for visual clarity) with the fitted hyperbola function (black dashed line). The glass transition temperature, $T_g = 211.7$ K, is identified as the centre of the hyperbola (red vertical line). The transition region (shaded area) is defined as the temperature range where the hyperbola derivative has converged to within 90% of its asymptotic values, spanning from $T_- = 192.8$ K to $T_+ = 230.6$ K. **B.** Temperature derivative of enthalpy, $dH/dT$, showing the sigmoidal transition between the glassy (green dashed line) and liquid (pink dashed line) asymptotic slopes. **C.** Bootstrap distribution of $T_g$ estimates obtained by refitting the hyperbola to 1000 synthetic datasets generated by adding Gaussian noise to the fitted model. The histogram shows the distribution of $T_g$ values, with the fitted normal distribution overlaid (blue line). The narrow distribution yields $T_g = 211.7 \pm 0.3$ K (uncertainty represents one standard deviation), demonstrating the robustness of the hyperbola method to statistical fluctuations in the enthalpy data. The well-defined, approximately Gaussian distribution validates the use of parametric bootstrap for uncertainty estimation and confirms that the $T_g$ estimate is statistically well-determined from the cooling trajectory.

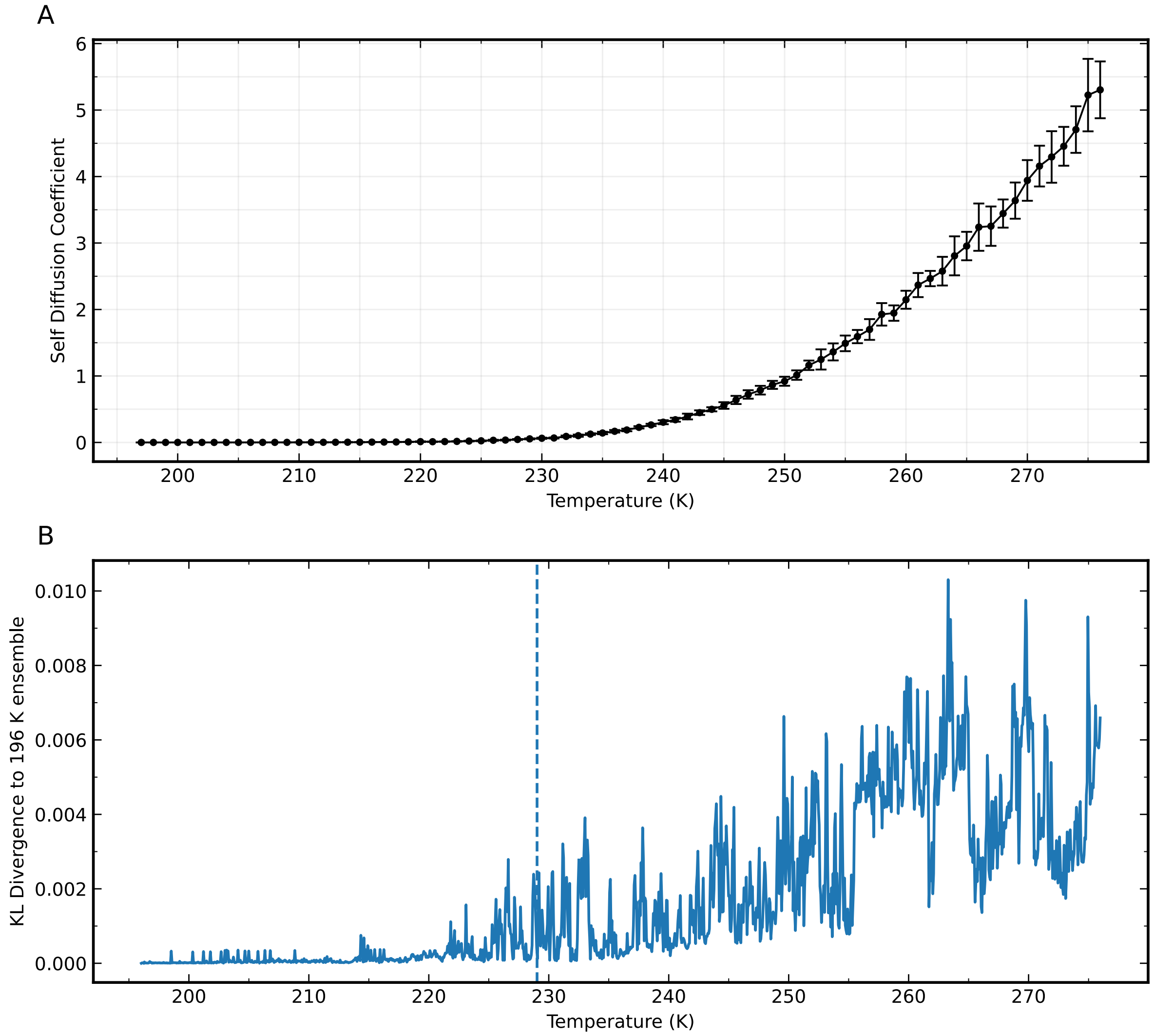


**Fig. S3 Temperature dependence of Trp-cage and solvent dynamics. A.** The self-diffusion coefficient of TIP4P/ICE model water as a function of temperature. Water self-diffusion coefficients were obtained from mean-squared displacement (MSD) [72, 73] analysis of the water molecules using the in-built MDAnalysis function [74, 75]. For each temperature, the three-dimensional Einstein MSD of water was evaluated over the final 5 ns of each temperature to ensure within the linear regime, and the MSD was fitted by linear least squares. Diffusion coefficients $D$ were then computed from the Einstein relation. $D = \frac{1}{2d}\frac{\mathrm{d}}{\mathrm{d}t}\left\langle |\mathbf{r}(t) - \mathbf{r}(0)|^2 \right\rangle$, with $(d = 3)$. Points represent the mean over 16 simulations and error bars show standard deviation. **B.** The KL Divergence of microstate distributions of Trp-cage as a function of temperature. Distributions are compared to the final frame. These 5341 pilot cooling simulations were carried out at a rate of 6.4 $\times 10^7$ K/s to 196 K.

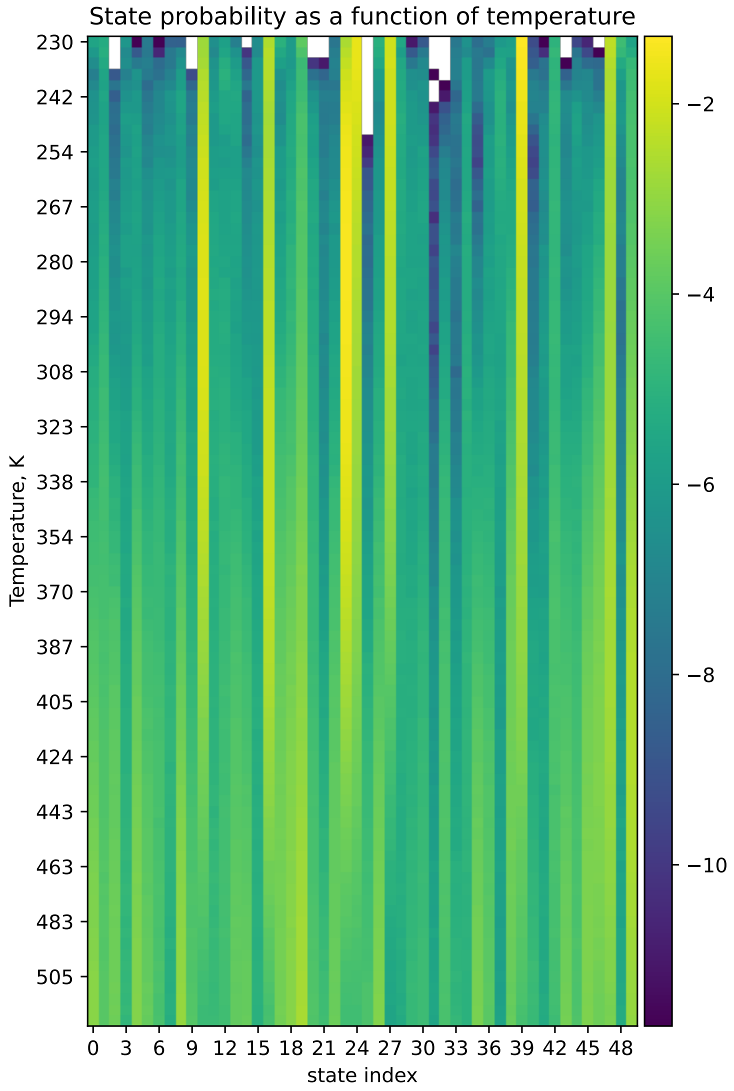


**Fig. S4 State probabilities from histogramming reassigned frames across each temperature from the TREX dataset**. Here we show the equilibrium probability at each temperature in our replica exchange ladder in terms of the state space we developed for our MSM at 277. As expected, probabilities are more uniform at higher temperatures, and become concentrated in a smaller number of more energetically favourable states at lower temperatures. Importantly, 10 states that are well populated in the 277 K ensemble are unobserved—there were no samples, indicated by a white value in the heat-map—at 230 K. Probabilities are in logscale.

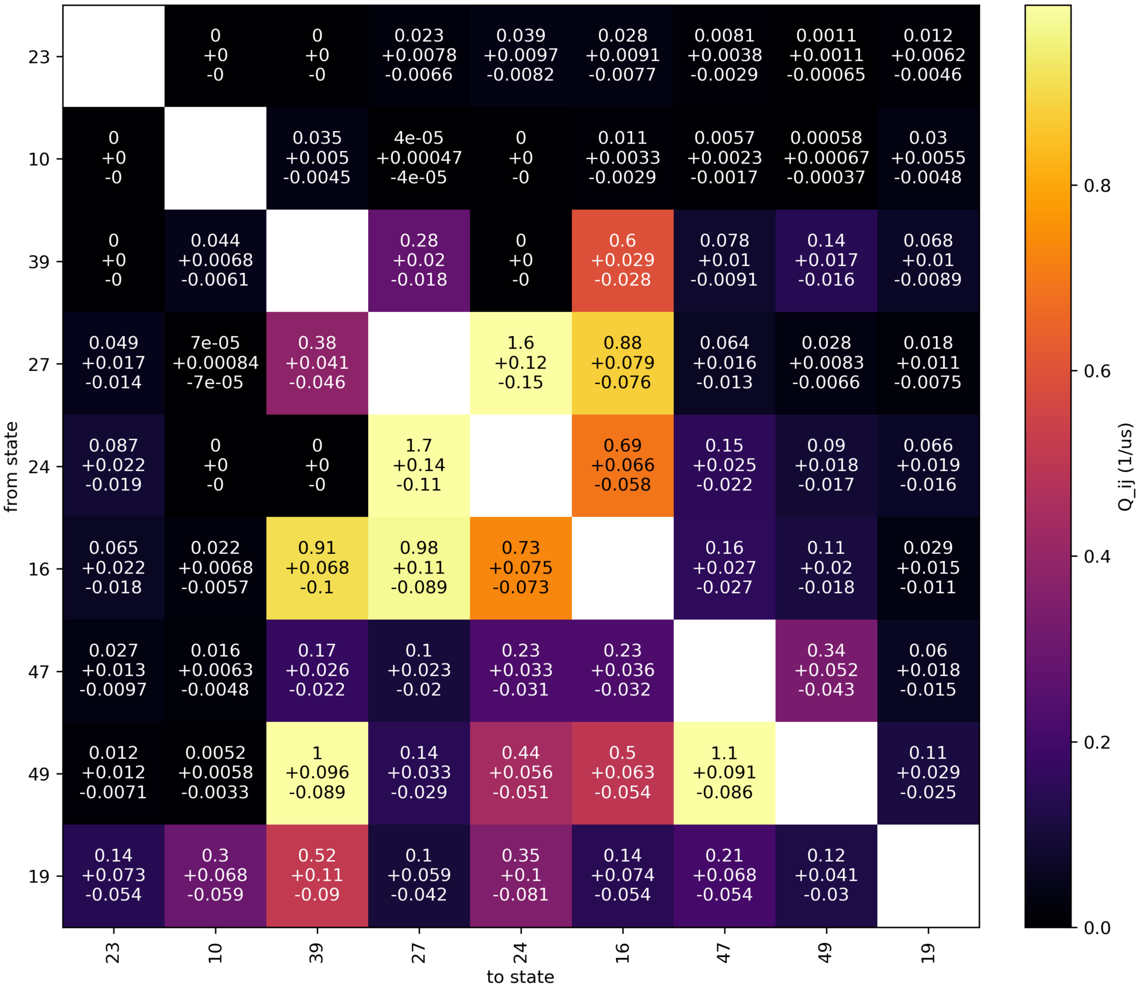


**Fig. S5 Transition rate matrix between the nine most populous microstates at 277 K.** State indices are provided on the X and Y axes, and the transition time from state i, indicated by the X-axis index, to state j, indicated by the Y-axis index, is provided as a colour associated to the colour bar on the left hand side in reciprocal µs. The text within the cells provides the median, plus and minus the 95% credibility interval for these quantities.

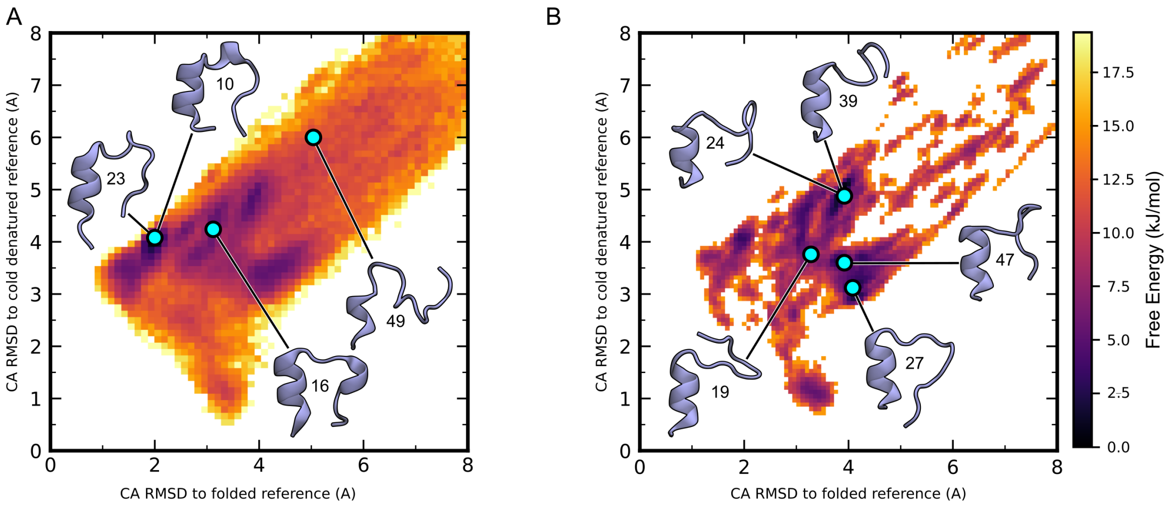


**Fig. S6 Microstate placement of Trp-cage conformers on 2D free energy landscape. A.** The free energy landscape of Trp-cage conformations at 277 K with representatives from microstates 10, 23, 16 and 49 annotated. These microstates have higher populations at 277 K versus 230 K. **B.** The free energy landscape of Trp-cage conformations at 230 K with representatives from microstates 24, 39, 27 and 47 annotated. These microstates have higher populations at 230 K versus 277 K. RMSD is calculated against the cluster centre of microstates 23 (folded) and 24 (cold-denatured). The positions of each state are at the bin to which each microstate had the highest density within this feature space. This means that annotated structures are not necessarily in a local well or close in geometric space. We prioritized using kinetic clustering to construct our MSM.

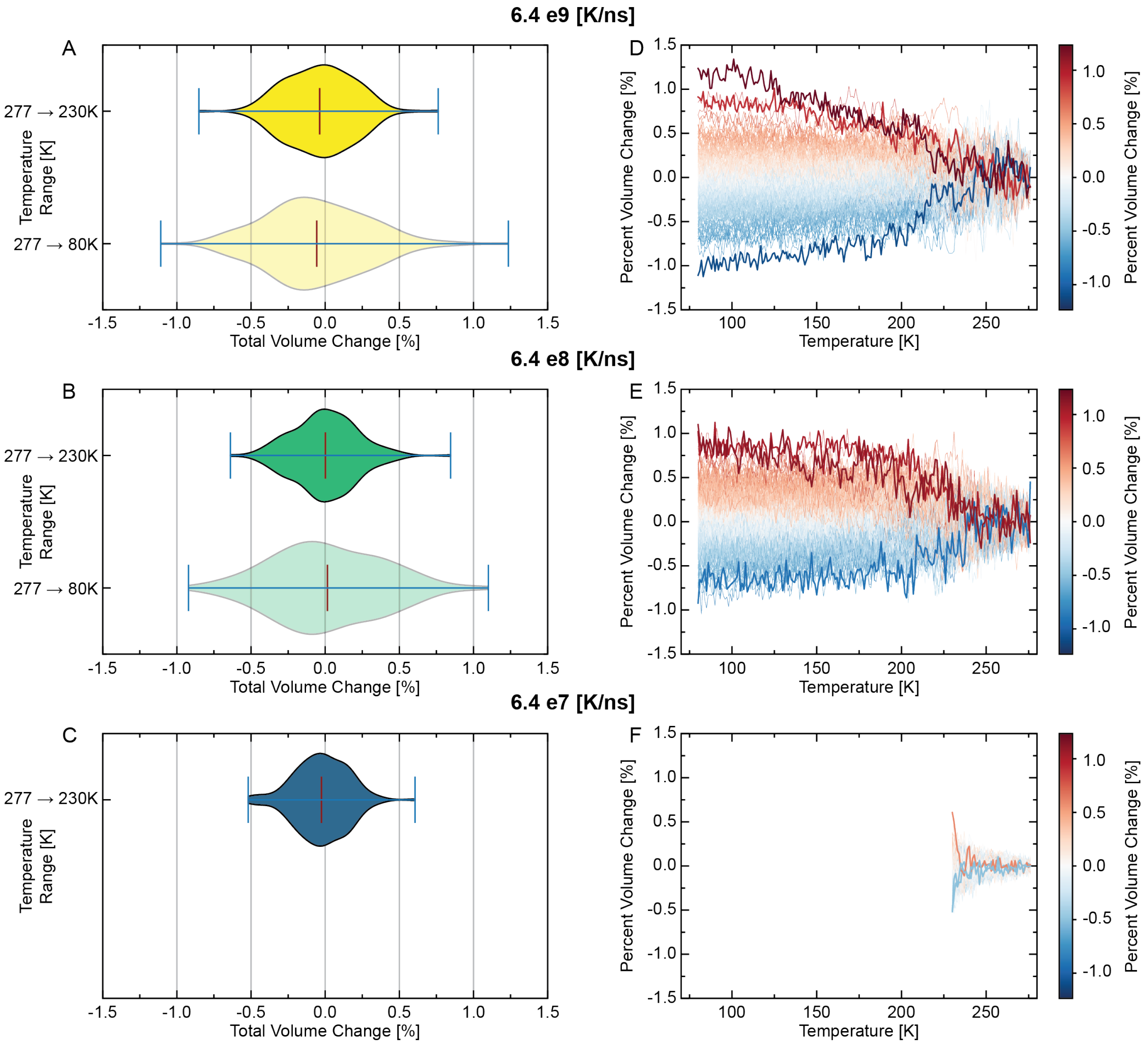


**Fig. S7 Cluster-resolved protein volume changes upon cooling.** Analysis of normalized protein volume changes across 200 conformational microstates for three cooling rates: $q_c = 6.4 \times 10^9$K/s (A, D; yellow), $6.4 \times 10^8$K/s (B, E; blue), and $6.4 \times 10^7$K/s (C, F;, blue). **Panels A, B, and C:** Violin plots showing the probability density distribution of volume changes relative to the high-temperature reference (276 K). For $q_c = 6.4 \times 10^9$ and $6.4 \times 10^8$K/s, two distributions are displayed: the full cooling range to 80 K (lighter, bottom violin) and a restricted range to 230 K (darker, top violin) for comparison with the slowest cooling rate. The violin width indicates the density of microstates at each volume change value. Black vertical lines indicate medians; red vertical lines indicate means. The distributions are tightly centered near zero, demonstrating minimal volume change across all cooling rates and temperature ranges. **Panels D, E, F:** Individual volume change trajectories for 200 clusters, coloured according to their final volume change using a diverging red-blue colormap (red = expansion, blue = contraction). The top three clusters with largest magnitude changes are highlighted with thicker lines. The narrow distribution of trajectories around zero change confirms that protein volume remains essentially constant during cooling. All data points represent averages over 10 independent replica simulations per cluster.

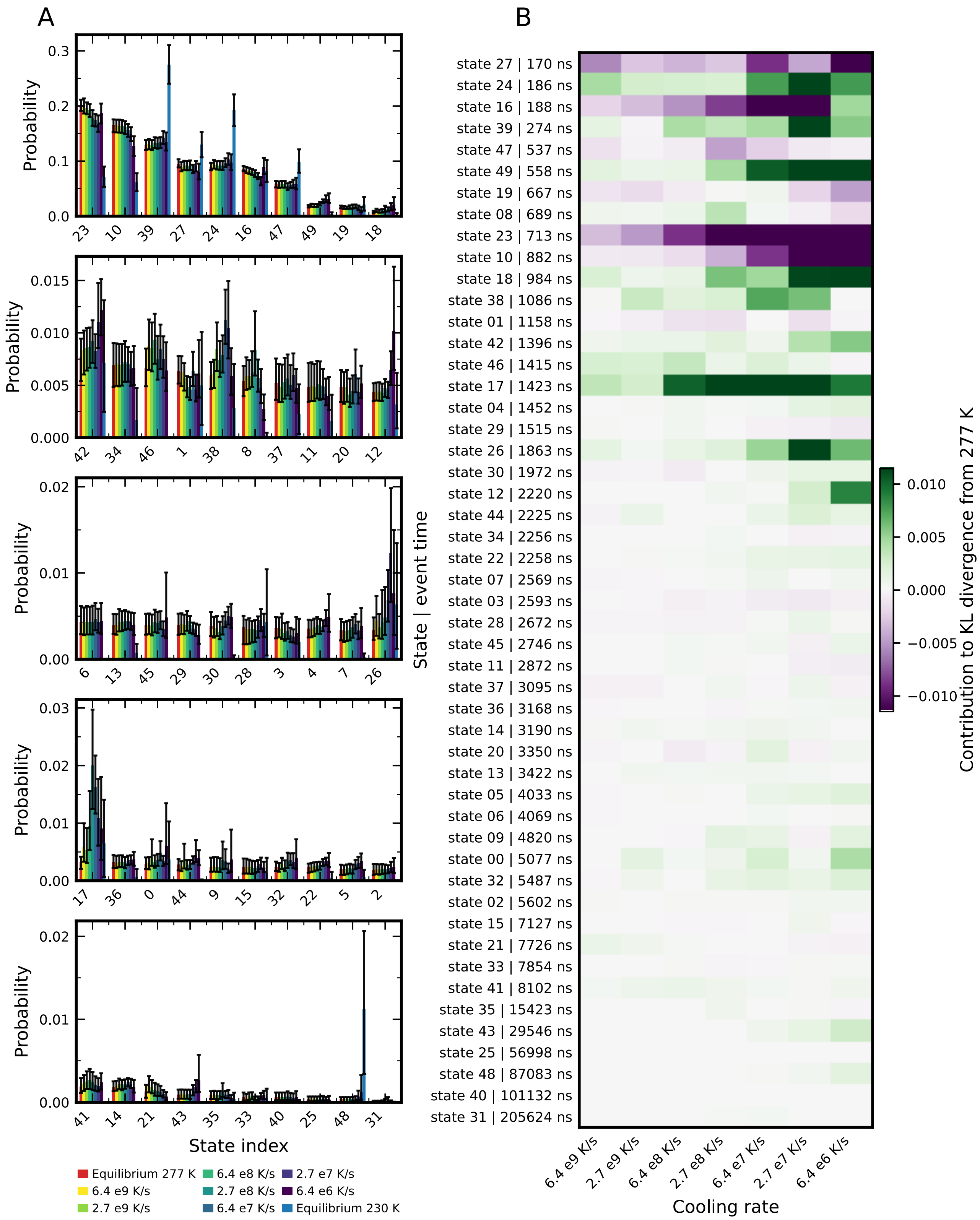


**Fig. S8 Behaviour of all 50 microstates in the MSM under cooling. A.** Evolution of microstates as a function of the cooling rate. Error bars represent 95% Credible Intervals. States are ordered by probability at 277 K at equilibrium. **B.** Heat map showing the relationship between state's event times, the cooling rate, and the change in that bin's contribution to the KL divergence between the equilibrium 277 K and cooled distributions, with the equilibrium as the reference.

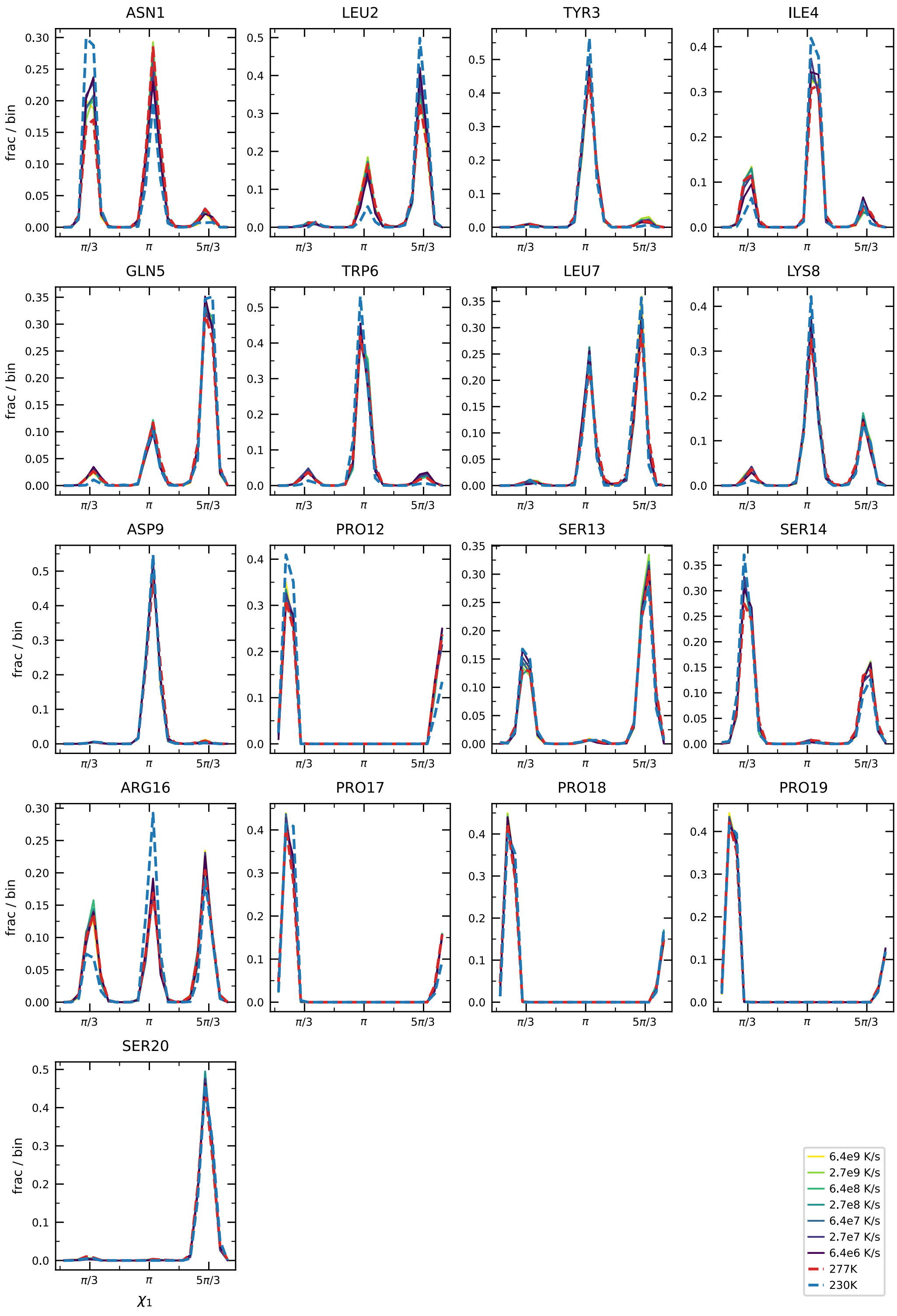


**Fig. S9** Cooled histograms of $\chi_1$ dihedrals for all 17 residues that have a side-chain. All angles are shifted by $\pi$ to translate the common rotamers a away from the periodic discontinuity. The dashed lines show the 277 K (red) and 230 K (blue) equilibrium histograms, with the viridis colour series showing the populations under cooling for each cooling speed from yellow (fastest) to purple (slowest), as in the main text. Some $\chi_1$ exhibit more heterogeneity than others, mostly corresponding to whether the sidechain can sample many conformations in the 277 K ensemble.

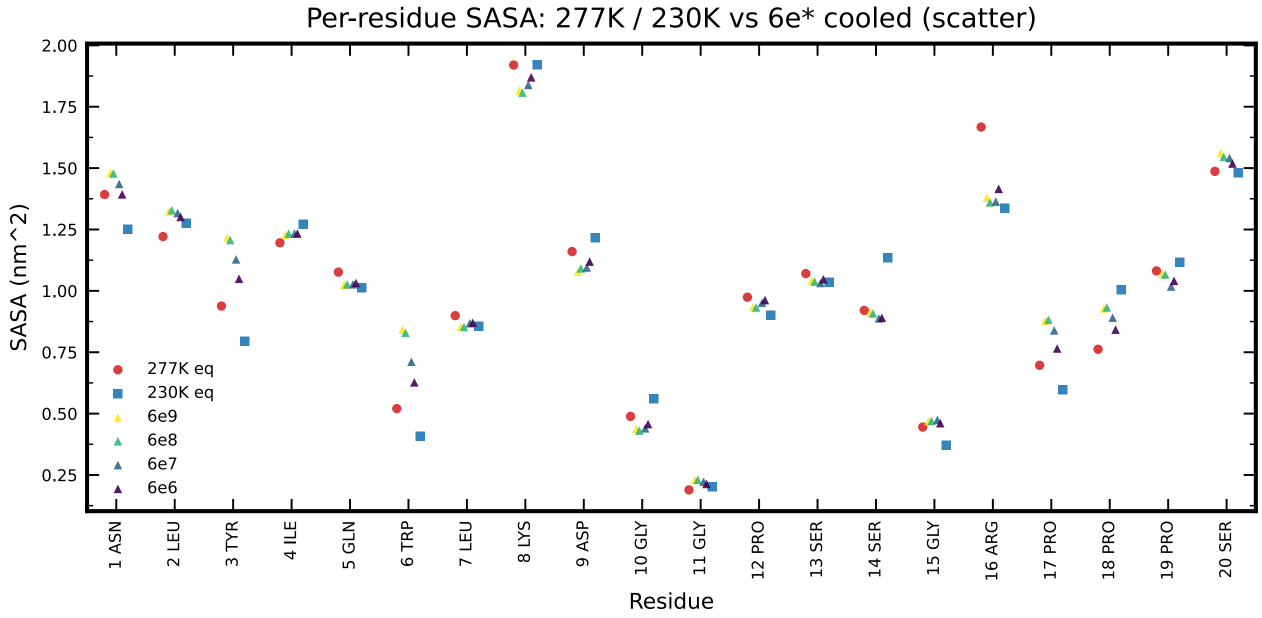


**Fig. S10** The population weighted Shrake-Rupley SASA per residue (as implemented in MDTraj[76], with default probe radius corresponding to water). The equilibrium 277 K ensemble is shown as red, the 230 K ensemble is in blue, and the cooled populations corresponding to the $6.4 \times 10^{X}$ cooling rates are shown as the viridis color series, matching the color coding used elsewhere in the text and supplement.

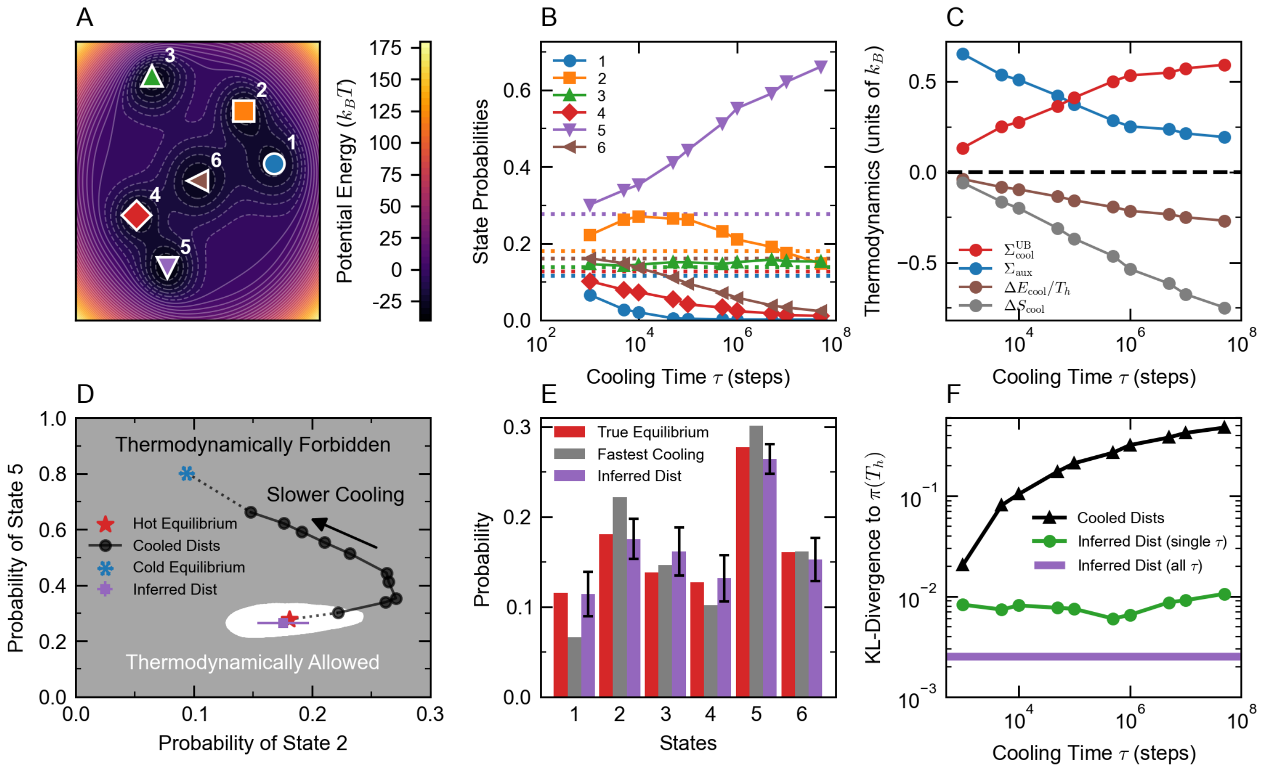


**Fig. S11** Illustration of the thermodynamic recovery method for a simple model consisting of a Brownian particle in a two-dimensional potential energy landscape. **A.** Two-dimensional potential energy landscape with 6 wells used to test the thermodynamic recover method. **B.** State probabilities as functions of cooling time. Horizontal dotted lines indicate hot equilibrium distribution. **C.** Thermodynamic quantities as functions of cooling time. **D.** Projection of hot and cold equilibrium and cooled distributions onto the space of probabilities of states 2 and 5. **E.** Histogram showing the true equilibrium distribution, cooled distribution with the fastest cooling time, and inferred distribution using all cooling times. **F.** KL-divergence from the true equilibrium distribution for each nonequilibrium cooled distribution (red), the inferred distribution using thermodynamic constraints with only a single cooling time (blue), and the inferred distribution using all cooling times (green).

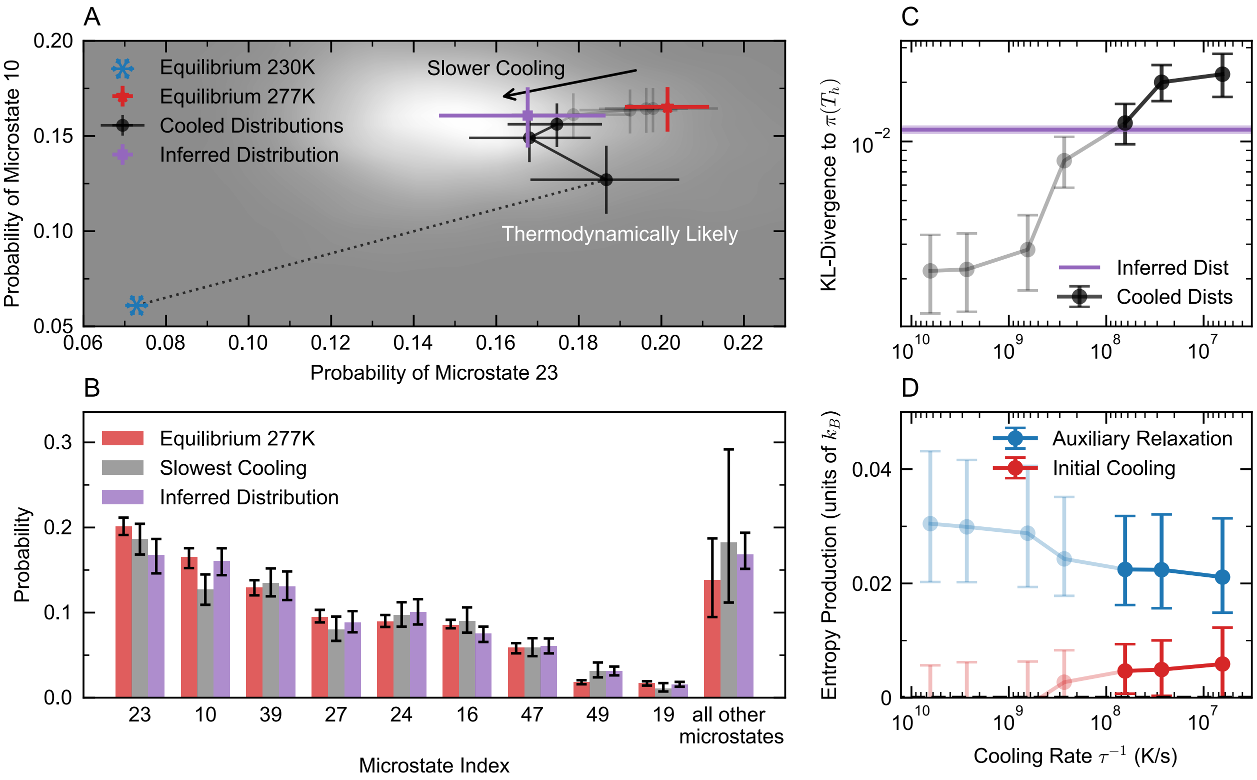


**Fig. S12 Using thermodynamic constraints to recover equilibrium ensemble from only the three slowest-cooled distributions A.** Projection of hot and cold equilibrium and cooled distributions onto the space of probabilities of states 2 and 4. Solid black points show the three slowest-cooled distributions, while transparent black points show the faster-cooled distributions (not used) **B.** Histogram showing the true equilibrium distribution, cooled distribution with the slowest cooling time, and inferred distribution. **C.** KL-divergence for each nonequilibrium cooled distribution from the hot equilibrium distribution (black), as well as between the hot equilibrium distribution and the inferred distribution (purple). Light purple band indicates error bars. **D.** Entropy production for the initial cooling process (UB, red) and auxiliary relaxation process (blue) as functions of the cooling rate $\tau^{-1}$, computed using the inferred state probabilities and entropies.

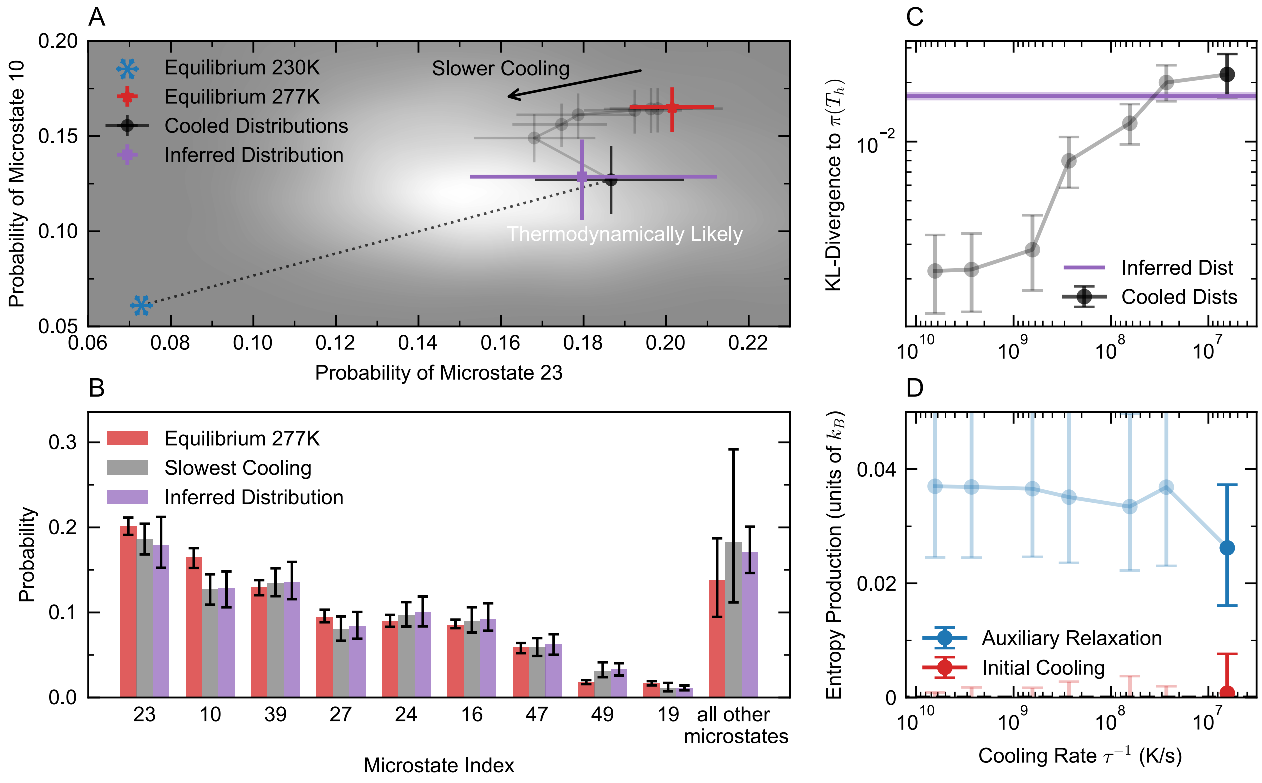


**Fig. S13 Using thermodynamic constraints to recover equilibrium ensemble from only the distribution cooled at the critical cooling rate. A.** Projection of hot and cold equilibrium and cooled distributions onto the space of probabilities of states 23 and 10. Solid black points show the slowest-cooled distribution, while transparent black points show the faster-cooled distributions (not used) **B.** Histogram showing the true equilibrium distribution, cooled distribution with the slowest cooling time, and inferred distribution. **C.** KL-divergence for each nonequilibrium cooled distribution from the hot equilibrium distribution (black), as well as between the hot equilibrium distribution and the inferred distribution (purple). Light purple band indicates error bars. **D.** Entropy production for the initial cooling process (UB, red) and auxiliary relaxation process (blue) as functions of the cooling rate $\tau^{-1}$, computed using the inferred state probabilities and entropies.

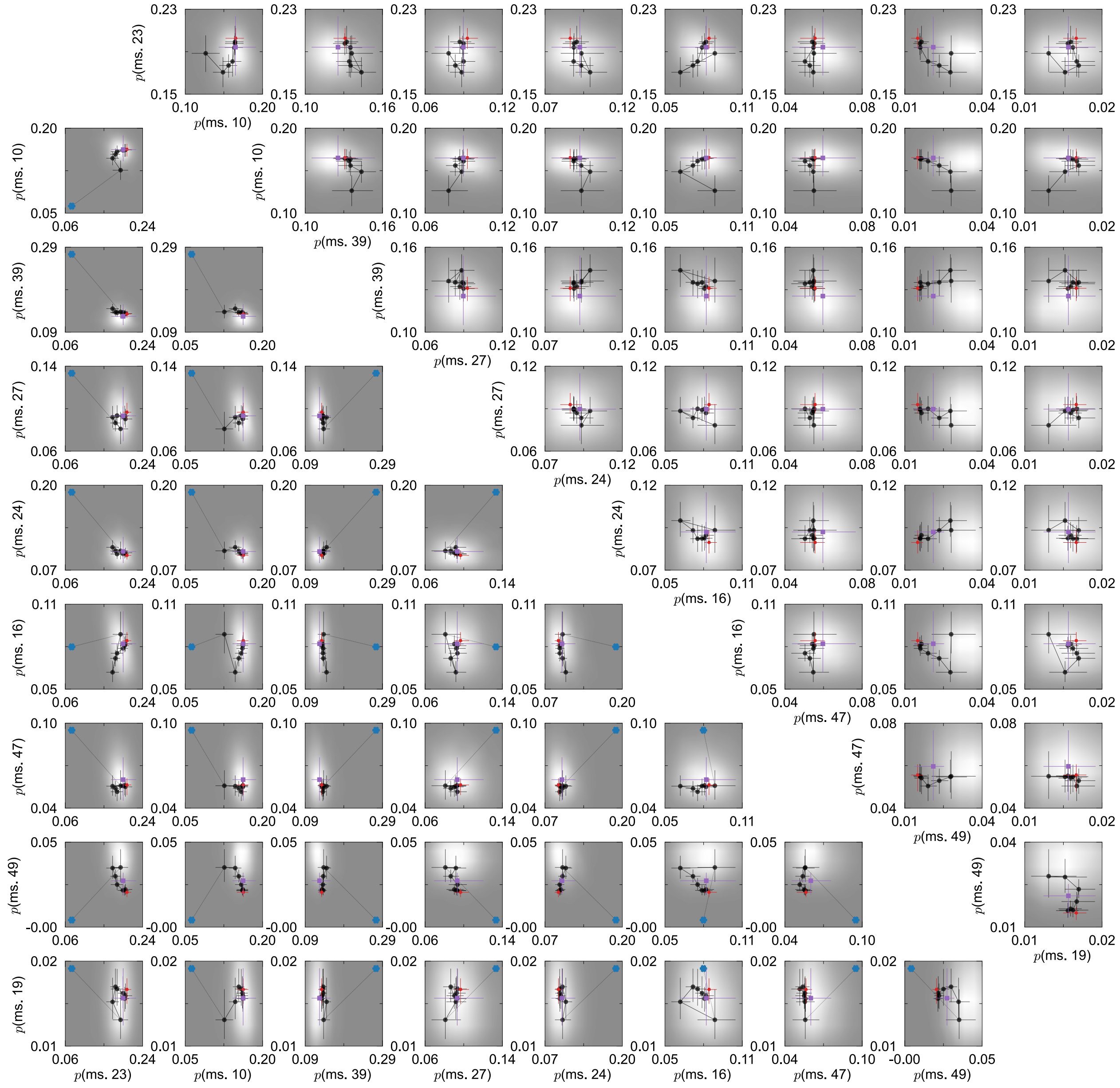


**Fig. S14** Every possible 2D projection, where both states have probability greater than 1%, of thermodynamic recovery for the microstate populations, both including (lower left) and excluding (upper right) the cold equilibrium distribution. Here the abbreviation "ms." refers to "microstate".

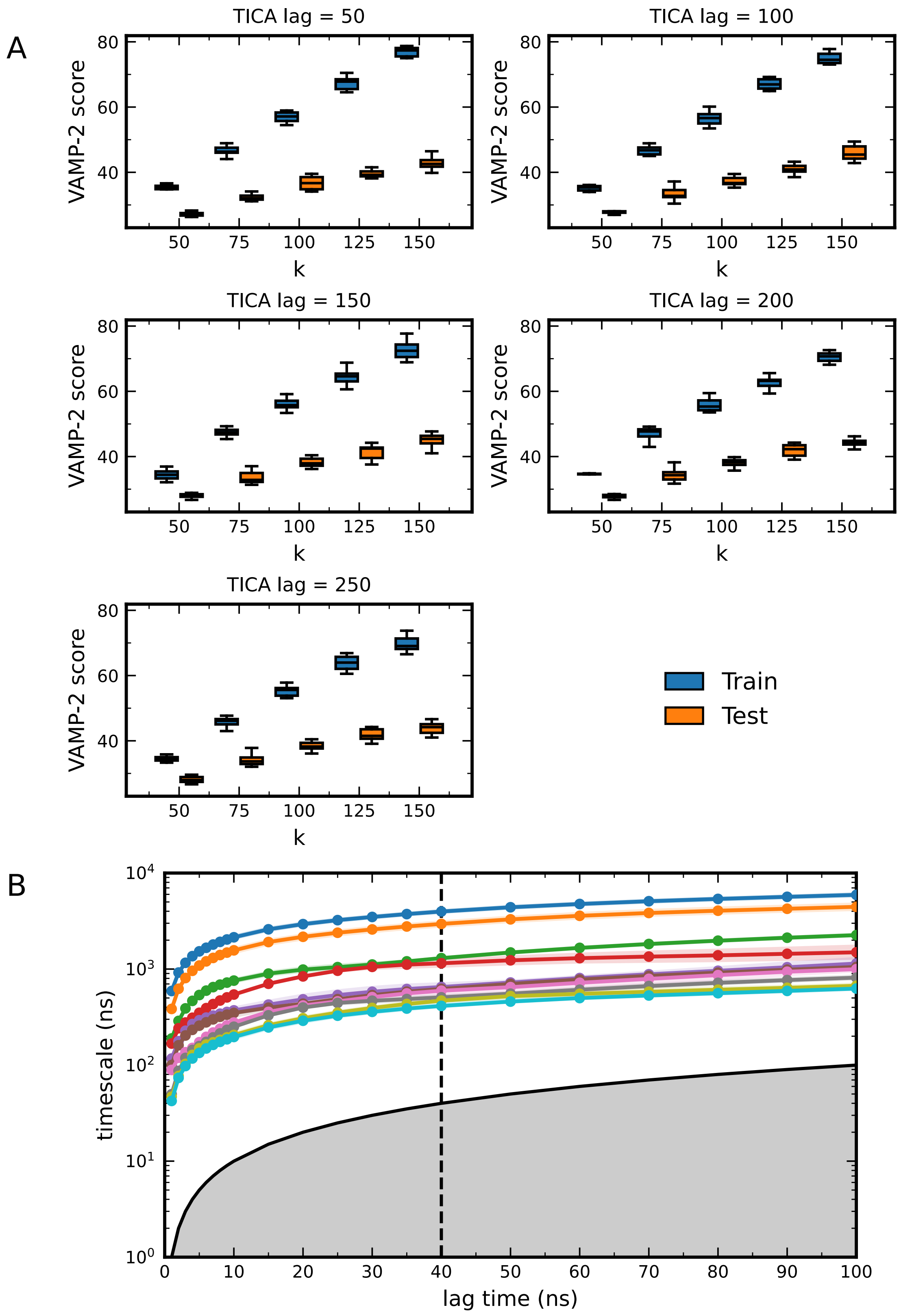


**Fig. S15 MSM parameter validation. A.** VAMP-2 scores across k clusters and TICA lags. **B.** Implied timescale plot of k = 50, TICA lag time = 25 ns. The lowest 10 processes are shown. Dashed black line at 40 ns indicates the chosen MSM lag for our models.

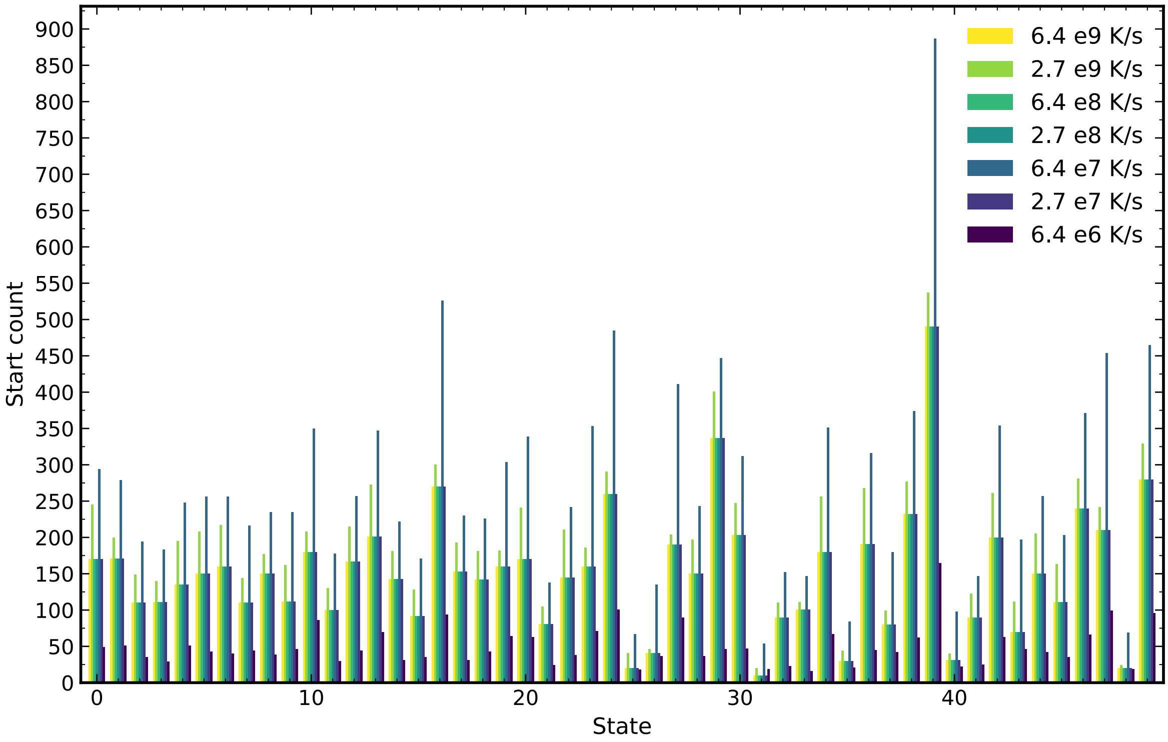


**Fig. S16  Start Counts of each microstate across cooling rates.**
